# A bioelectrical phase transition patterns the first beats of a vertebrate heart

**DOI:** 10.1101/2022.12.06.519309

**Authors:** Bill Z. Jia, Yitong Qi, J. David Wong-Campos, Sean G. Megason, Adam E. Cohen

**Affiliations:** Department of Chemistry and Chemical Biology, Harvard University, MA 02138; Department of Systems Biology, Blavatnik Institute, Harvard Medical School, MA 02115; Systems, Synthetic and Quantitative Biology Ph.D. Program, Harvard University; Department of Physics, Harvard University, MA 02138

## Abstract

The heart is among the first organs to function in vertebrate development, but its transition from silent to beating has not been directly characterized. Using all-optical electrophysiology, we captured the very first zebrafish heartbeat and analyzed the development of cardiac excitability around this singular event. The first beats appeared suddenly and propagated coherently across the primordial heart. Targeted optogenetic perturbations mapped the development of excitability and conduction before and after the first heartbeats. Measured bioelectrical dynamics support a noisy saddle-node on invariant circle (SNIC) bifurcation as the critical phase transition that starts the heart. Simple models of this bifurcation quantitatively capture cardiac dynamics in space and time through early development, including coherent beating before transcriptional specification of pacemakers. Our work shows how gradual and largely asynchronous development of single-cell bioelectrical properties produces a stereotyped and robust tissue-scale transition from quiescence to coordinated beating.

## Main Text

Cardiac activity during early embryonic development has been documented for over two thousand years (*1, 2*). However, *in vivo* observations of physiological function in the heart primordium have been limited in spatiotemporal resolution and sample size (*3*–*7*). This scarcity of data means that the bioelectrical mechanisms for the emergence of organized cardiac function are still poorly understood. We asked: how does the heart go from silent to regular beating? What are the intermediate activity states, and how do these states emerge from the ensemble of single-cell developmental trajectories (*8*)? Here, we applied all-optical electrophysiology (*9, 10*) to observe the first beats of developing zebrafish hearts and to assess the electrical excitability and connectivity patterns underlying these dynamics.

A first heartbeat is a once-in-a-lifetime event. We sought to capture this event via calcium imaging as zebrafish embryos developed from 18 – 22 hours post fertilization (hpf; Fig. 1A). During this period, the bilateral cardiac progenitor populations converge and form the heart cone, and early heartbeats have been reported at homologous developmental stages in the chick, rat, and mouse (*3*–*7*). The progenitors are undergoing active differentiation at this stage, so we ubiquitously expressed the calcium sensor jGCaMP7f (*11*) to capture activity that might be missed by tissue-specific expression. To acquire robust statistics, we combined a system for wide-area all-optical electrophysiology (*12*) with a custom agarose mold for live imaging of up to 18 embryos concurrently (Fig. 1B; Fig. S1, Movie S1, S2).

**Figure 1.**
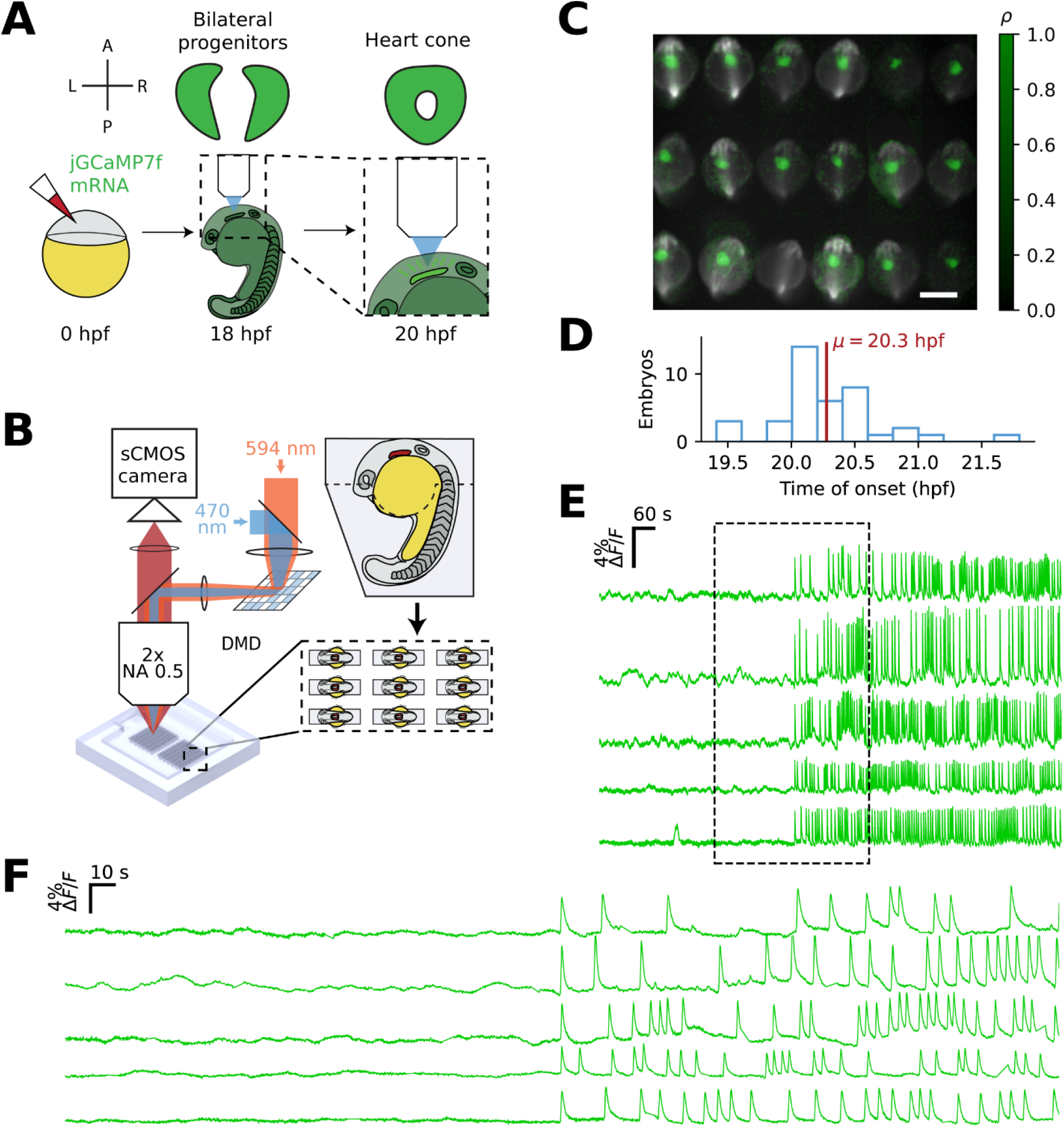
Multiplexed time-lapse calcium imaging captures the first heartbeats of zebrafish embryos. (**A**) Injection of mRNA encoding jGCaMP7f for whole-embryo Ca^2+^ imaging during zebrafish heart cone formation. (**B**) Low-magnification microscopy platform and array mount for multiplexed live imaging and all-optical electrophysiology. (**C**) Pearson correlation (green) of individual pixel time traces to putative cardiac activity. Activity maps are overlaid on grayscale images of baseline fluorescence. Scale bar, 500 µm. (**D**) Histogram of time of onset of the first cardiac calcium spike (*n* = 39 embryos, 3 experiments). (**E**) jGCaMP7f ΔF/F measurements of representative hearts aligned by the first calcium spike show abrupt onset of cardiac activity. (**F**) Zoomed-in traces from (**E**).

Within each embryo, the primordial heart showed an abrupt transition from quiescence to calcium spiking with a stereotyped waveform. These transitions occurred in a tight developmental window (20.3 ± 0.4 hpf, mean ± SD, N = 39 fish; Fig. 1C – E, Methods). Experiments in fish expressing the red calcium sensor FR-GECO1c (*13*) showed that these events colocalized with *nkx2*.*5:ZsYellow* (Fig. S2), a marker of the heart primordium at this stage of development (*14*). Consistent with previous reports, rhythmic calcium dynamics were present before mechanical contractions could be detected (typically 21-22 hpf) using brightfield microscopy (Movies S3 and S4) (*4, 5, 7*). We refer to the first large-scale calcium transient as the first heartbeat.

We next asked whether the first heartbeat engaged the whole tissue or was confined to a subset of cells. Prior to the first heartbeat, we observed occasional single-cell calcium transients, as has been reported previously (*7*). These events were infrequent (0.2 – 0.7 per minute), long-lived (3.40 ± 2.09 s, mean ± SD, n = 19 events, 3 animals), lacked a sharp peak, and were asynchronous between cells, making them qualitatively distinct from the first heartbeats (Fig. 2A, Fig. S3). The first beat usually engaged most of the heart primordium, engaging > 4000 μm^2^ of tissue, compared to a typical cell size of 82 μm^2^ (Fig. S3 – S4; Movie S5). However, the first beat occasionally (3/39) preceded the electrical fusion of the two progenitor fields and only propagated through half of the heart (Fig. 2A, Fig. S5). In such cases, the beat engaged the contralateral side within 30 minutes. During the hour after the first beat, the spatial extent of the spikes and the spatially averaged spike amplitude only increased slightly (area: 35 ± 47% increase, amplitude: 22 ± 27% increase, mean ± SD, *N* = 39 fish) and spike width only decreased slightly (−24±10% decrease, mean ± SD, *N* = 39 fish), (Fig. 2, B – D; Fig. S6). Therefore, the initiation of the heartbeat was a step change from sparse and slow single-cell transients to sharp, tissue-scale spikes that remained relatively stable over the following hour of development.

**Figure 2.**
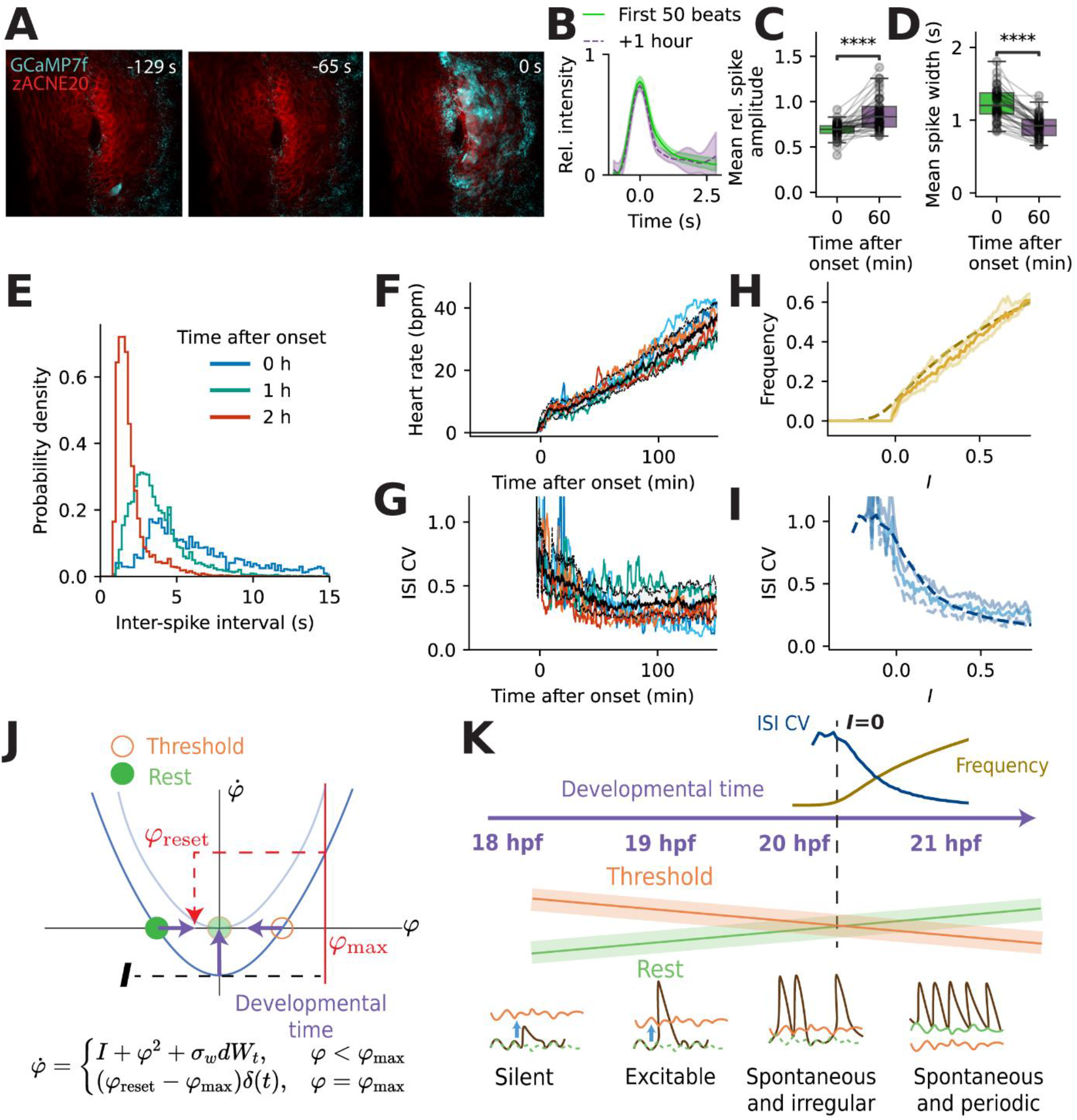
The heartbeat undergoes coordinated initiation via a noisy SNIC bifurcation. (**A**) Images showing isolated single-cell calcium transients followed by a large-scale coherent event at *t* = 0. Scale bar, 50 µm. (**B**) Example spike-triggered averages for individual embryos at onset and 1 hour post onset (normalized to the largest amplitude in the first 50 beats). (**C**) Relative transient amplitude increased by 22±27% (mean ± SD) in the first hour after onset. Paired t-test, **** *p* = 6.3e-6. (**D**) Transient full-width half-maximum decreased by 24±10% (mean ± SD) in the first hour after onset. Paired t-test, **** *p* = 2.4e-15, fold change of median 0.76. (**E**) Distribution of inter-spike intervals (ISIs) in selected 10-minute windows relative to the first heartbeat. (**F**) Beat rates increased in a stereotyped trajectory. (**G**) Interspike interval coefficient of variation (ISI CV) decreased in a stereotyped trajectory. (F – G) Black line: median; dotted lines and shading: interquartile range; colored lines: representative individual traces. (**H** and **I**) Experimental frequency and CV jointly fit to the quadratic integrate-and-fire (QIF) model (dashed line, Supplementary Text, Fig. S4) with only linear scalings of drive current to developmental time. Shaded region is ±95% CI. (B – G) Same individuals as Fig. 1D, *n* = 39 embryos, 3 experiments. (**J**) Phase portrait of the QIF model as it undergoes a SNIC bifurcation. When the spiking variable *φ* is to the left of the unstable saddle (orange), it decays to the stable node (green). When *φ* is to the right of the saddle, it grows to *φ*_*max*_ and then is moved back instantaneously to *φ*_*reset*_. (**K**) Emergence of spiking regimes in the early heartbeat driven by a noisy phase transition.

We then investigated how the heartbeat became periodic. The first beats were infrequent and irregularly spaced, and the inter-spike interval (ISI) became shorter and more uniform as the heart developed (Fig. 2, E). The heart rate increased from 7.1 ± 3.3 beats per minute (bpm) at 10 minutes post onset (mpo) to 29.6 ± 7.9 bpm at 120 mpo (mean ± SD, *N* = 39 fish) (Fig. 2F). We quantified the heart rate variability by the coefficient of variation (CV: standard deviation/mean) of the ISI. This ISI CV decreased from 0.62 (median; interquartile range, IQR 0.44-0.83) at 10 mpo (ISI CV = 1 corresponds to a Poisson process) to 0.37 (median; IQR 0.26-0.52) at 120 mpo (Fig. 2G).

The mean heart rate and ISI CV followed stereotyped trajectories when aligned temporally to the first beat, suggesting a characteristic transition underlying the dynamics (Fig. 2, F and G). We thus sought to understand the nature of the transition from quiescence to beating. This transition can be described as a codimension-1 bifurcation, i.e. a step change in dynamics driven by a continuous change in a control parameter. There are just four classes of codimension-1 bifurcations into an oscillatory state (*15*). We hypothesized that by assigning the transition to one of these classes, we could formulate a model that reproduced the observed dynamics and possibly even predict the response to external perturbations.

We compared our experimental results to simulated excitable-to-oscillatory transitions of each bifurcation class, using the Morris-Lecar action potential model with the addition of noise to recapitulate the ISI CV (Supplementary Text, Fig. S7) (*16*). Only the saddle-node on invariant circle (SNIC) bifurcation captured the statistics of our data. All SNIC bifurcation models have similar dynamics near the bifurcation point (*17*), so we decided to test the quadratic integrate-and-fire (QIF) model, which is the simplest member of the class (Supplementary Text, (*15, 18*)). The noisy QIF model comprises just a “spiking” variable, *φ*, and two free parameters, the “injected current”, *I*, and the noise power, *σ*_*w*_^*2*^. Strikingly, within a range of values of *σ*_*w*_^*2*^, the noisy QIF model captured the evolution of both mean beat frequency and ISI CV with just a linear scaling of *I* to developmental time (Fig. 2, H and I; Fig. S8). Further, the noisy QIF model predicts a relatively constant spike amplitude as *I* grows near the bifurcation point, and an absence of bistability between silent and spiking states, both consistent with our data. In summary, the periodic heartbeat emerges through a phase transition-like SNIC bifurcation, bridged by a regime of noisy spontaneous spiking (Fig. 2, J and K).

A key prediction of the SNIC bifurcation model is that the heart should be excitable before it becomes spontaneously active. To test this prediction, we co-expressed the far-red calcium sensor FR-GECO1c (*13*) and the channelrhodopsin CoChR (*19*) (Fig. 3A), either by mRNA injection at the one-cell stage, or in a transgenic line with a novel chimeric promoter (*zACNE20-myl7*) engineered to drive early and strong expression in the developing heart (Fig. S9, (*20*)). Using our multiplexing platform, we applied periodic impulses of blue light targeted to the whole heart, and measured the probability of evoking a calcium transient as a function of developmental time (Fig. 3B, Movie S2). We observed all-or-none responses to individual stimuli, indicative of coherent tissue-wide responses similar to later spontaneous heartbeats. The heart became optogenetically excitable in a window 90 minutes before the first spontaneous beats (Fig. 3, B – D), qualitatively matching our expectation from theory. Within individual embryos, the response rate to optogenetic stimuli increased from 0% to close to 100% in typically less than 5 minutes.

**Figure 3.**
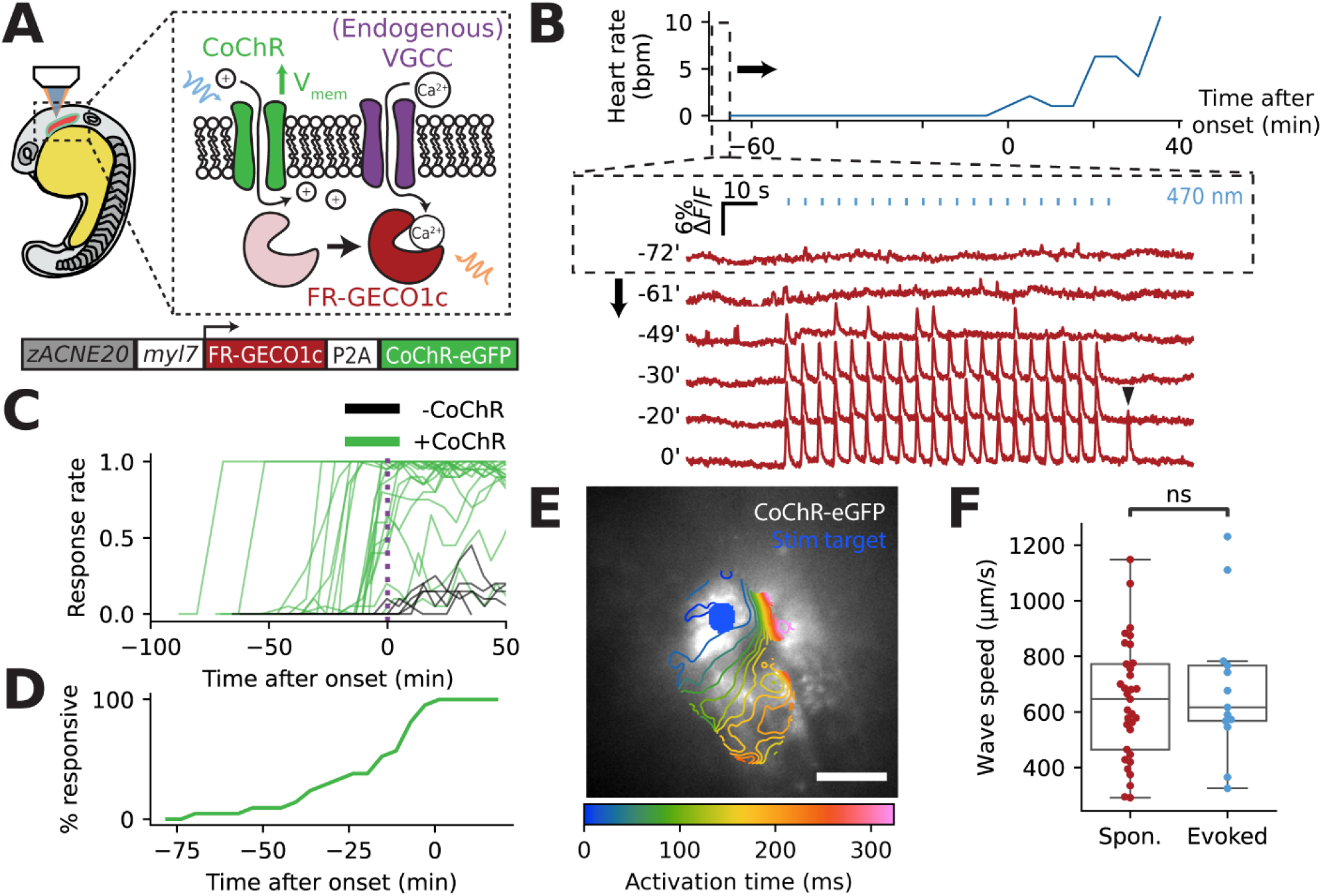
The heart primordium is excitable and electrically coupled before the first heartbeat. (**A**) Genetic construct for simultaneous optogenetic stimulation and calcium imaging, driven by a chimeric zACNE20-myl7 promoter for early expression in the heart. (**B**) Repeated testing of response to optogenetic stimulation (twenty 25 ms pulses at a period of 5 seconds) showed increasing excitability before onset of spontaneous activity. Black triangle indicates the first detected spontaneous beat. (**C**) Response rate to optogenetic stimuli increased from 0% to 100% for most embryos before onset of the spontaneous heartbeat. Response rate for embryos not expressing CoChR was a result of spontaneous transients randomly aligning with blue light pulses. (**D**) Most embryos gained excitability in the hour before onset of spontaneous activity, as quantified by the fraction of the population displaying a greater than 50% response rate. (C – D) *n* = 5 embryos -CoChR, *n* = 21 embryos +CoChR, two experiments. (**E**) Prior to onset of spontaneous heartbeat, spatially targeted CoChR stimulation elicited propagating calcium waves. Isochronal lines for time at which each pixel reached a maximum in d(ΔF/F)/dt. Scale bar, 50 µm. (**F**) Median propagation speed was ∼625 µm/s in waves evoked before onset of spontaneous activity and in early spontaneous waves (< 30 minutes after the first beat). Evoked, thirteen 45-second recordings, 7 embryos, 2 experiments. Spontaneous, ninety 30-second recordings, 12 embryos, 3 experiments. Mann-Whitney-Wilcoxon two-sided test, n.s. *p* = 0.46.

During the epoch where the heart was excitable but not yet spontaneously active, we targeted stimuli to small sub-regions of the heart. These stimuli elicited propagating calcium waves that initiated at the stimulus location (Fig. 3E; Movie S6). Therefore, cells were electrically coupled before the initiation of the heartbeat and could be excited not only by external stimuli but also by their neighbors. Wave propagation speeds were similar between precocious optically evoked waves and early spontaneous activity (median 617 µm/s vs. 646 µm/s, n.s., N = 7 evoked, 12 spontaneous) (Fig. 3F). These speeds were an order of magnitude faster than reported values for calcium waves driven by diffusion of small molecules (*21*), suggesting that propagation was mediated by gap junctional coupling of membrane potential.

Morpholino knockdown of the *cacna1c* gene encoding the L-type voltage-gated calcium channel *α*1c subunit abolished spontaneous (N = 11 embryos) and evoked (N = 14 embryos) calcium transients (Fig. S10). Thus the first beats comprised regenerative action potentials driven primarily by a calcium current, consistent with prior observations in the 36 hpf zebrafish heart(*22*). The presence of both electrical excitability and gap junctional coupling thus primes the heart for a tissue-wide response to the first spontaneous action potential, explaining the sudden transition from silence to large-area coherent heartbeats.

Spontaneous activity is widespread in the embryonic heart and this activity gradually becomes localized to molecularly distinct pacemaker regions during development (*23*–*26*). Reports of the initial pacemaking locus of initiation (LOI) are conflicting (*3*–*5*), and the mechanisms determining its location in the early heart are unknown. We performed jGCaMP7f imaging at 100 Hz to generate activation maps of the spontaneous heartbeats at 20-22 hpf (Fig. 4A). We identified the location with the earliest upstroke of each calcium transient as the spontaneous LOI (Methods). The LOI was most frequently in the anterior-left quadrant of the heart cone (38/71 observations across 10 embryos) and was relatively stable over time, although some motion and variability between samples were observed (Fig. 4B). Large and sudden displacements in the LOI position occurred in a few embryos.

**Figure 4.**
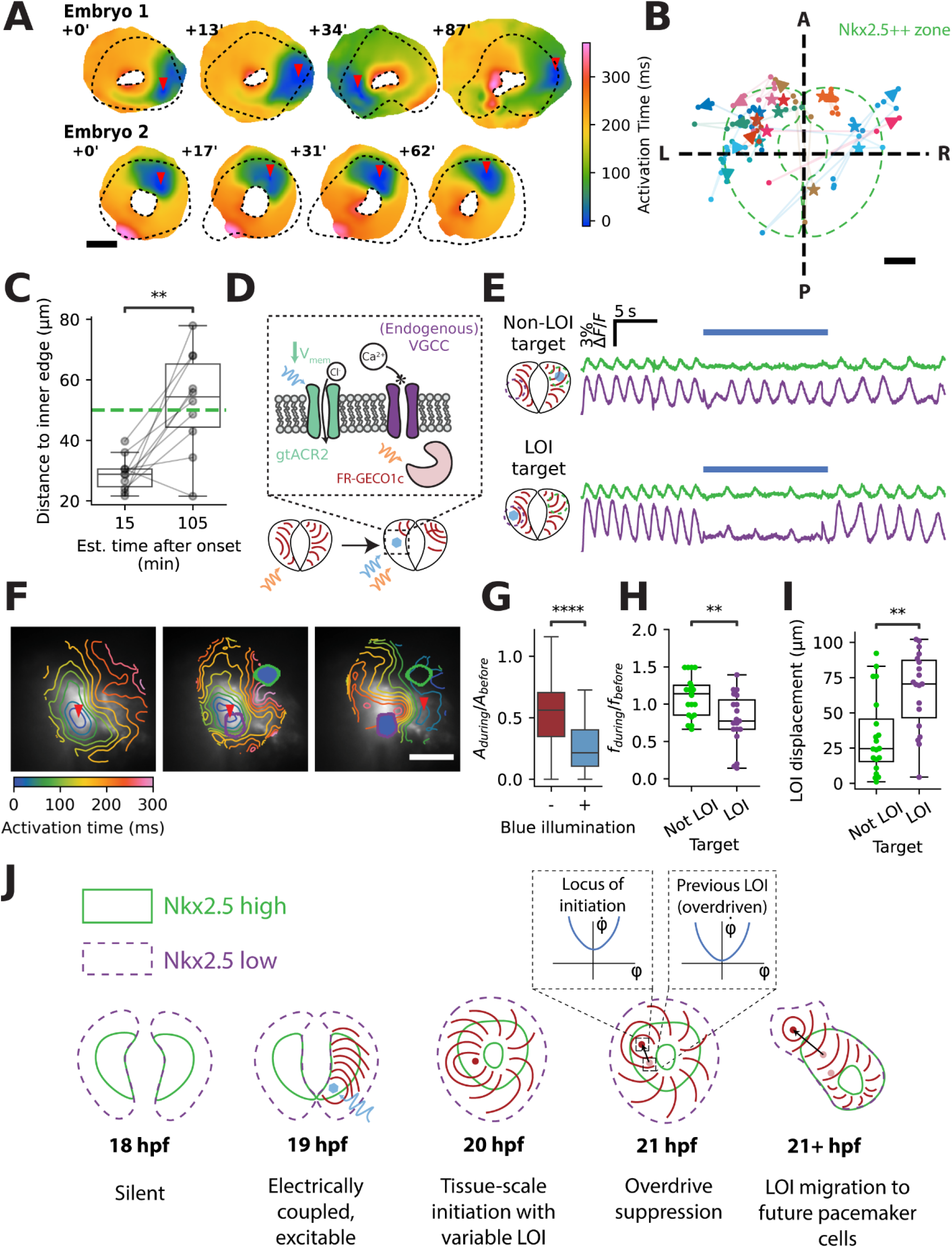
Wave geometry of the early heartbeat is set by competition between spontaneously firing cells. (**A**) Activation maps for spontaneous activity of two embryos across time. Most embryos had a relatively stable locus of initiation (LOI, red arrow) (bottom), while some had substantial movement of the LOI (top). Dashed black line indicates the strongly Nkx2.5-positive (Nkx2.5++) region. Scale bar, 50 µm. (**B**) LOI trajectories for individual embryos show primarily anterior-left localization. Star indicates initial measurement, arrowhead indicates final measurement. Time between initial and final measurements was 45-240 minutes. Imaging intervals varied from 5.5 to 30.5 minutes (mean 14.6 minutes). Green dashed line indicates the approximate extent of the Nkx2.5++ region. A: anterior, P: posterior, L: left, R: right. Scale bar, 25 µm. (**C**) LOIs drifted outwards from the inner edge of the heart cone as development progressed. Estimated time after onset based on heart rate. Paired t-test, ** *p* < 0.01. (B – C) *n* = 12 embryos, three experiments. (**D**) Expression of gtACR2 enabled localized optogenetic hyperpolarization which prevented activation of voltage-gated calcium channels (VGCC). (**E**) Example FR-GECO1c ΔF/F traces showing localized silencing. Targeting a non-LOI region of interest (ROI) did not slow the heartbeat, but targeting the LOI slowed the heartbeat globally. Purple: fluorescence at spontaneous LOI; green: fluorescence at non-LOI region. Blue circle: optogenetic stimulus target. (**F**) Activation maps for the experiment in (E). Red triangles indicate LOI. Purple and green circles indicate region of interest whose fluorescence is plotted in (E). Left: endogenous activity. Center: targeted silencing (filled blue spot) of a non-LOI ROI. Right: targeted silencing of the endogenous LOI caused a new LOI to emerge. Scale bar, 50 µm. (**G**) Focal optogenetic silencing decreased the amplitude (A) of the calcium transients more in the directly illuminated region (+) than in regions distal from the optogenetic silencing (-). (**H**) Optogenetic silencing of the LOI reduced the heart rate more than did optogenetic silencing of a non-LOI region. (**I**) Optogenetic silencing of the LOI led to larger LOI displacement compared to silencing of a non-LOI region. (G – I) 39 observations in 13 embryos across 5 experiments. Mann-Whitney-Wilcoxon two-sided test, ** *p* < 0.01, **** *p* <10^−4^. (**J**) Establishment of spatial patterning in the early heartbeat. When the heart becomes electrically excitable (∼19 hpf), it also supports wave propagation. This means that emergence of spontaneity at a single point in the Nkx2.5++ heart cone induces tissue-wide activity. The LOI is determined by the region with the fastest natural frequency (Fig. 2I). Eventually the LOI migrates to an Isl1+/Nkx2.5-population in the inflow tract.

The early LOI tended to reside in the proximal and strongly Nkx2.5+ region of the heart cone (< 50 µm away from the inner edge of the heart cone), and drifted outwards as development progressed (Fig. 4, B and C; Fig. S4). This trajectory contrasts with the concurrent specification of a reported pacemaker lineage in a distal (∼100 µm outwards) and sparse Isl1+/TCF+ population undergoing downregulation of Nkx2.5 (*27*). Our observation is, however, consistent with previous descriptions of contractions beginning first in the future conoventricular region of the heart (*3, 28*). Therefore, the early LOI is spatially distinct from the cells that ultimately become pacemakers.

The SNIC model predicts that the spontaneous frequency depends strongly on the bifurcation parameter. Within a tissue where there are gradients and possibly noise in developmental trajectories, one would expect a diversity of spontaneous frequencies. How, then, does the tissue produce coherent beats that emerge from a relatively stable LOI? We hypothesized that gap junctional coupling led to phase locking to the highest frequency oscillator via overdrive suppression. To test this hypothesis, we used the hyperpolarizing anion channelrhodopsin gtACR2 (*29*) and targeted illumination to selectively silence different regions of the heart (Fig. 4, D–G, Movie S7), while simultaneously monitoring calcium with FR-GECO1c. Silencing of the LOI caused a drop in spontaneous frequency and the emergence of a new LOI in a different spot, whereas silencing non-LOI regions caused significantly smaller frequency shifts and little LOI displacement (Fig. 4, H and I, Movie S7). These observations indicate the presence of multiple potential pacemakers, with the LOI set by the one with the fastest natural frequency. This mechanism resembles the ”overdrive suppression” that sets pacemaker position in the adult cardiac conduction system (*30*).

We further tested the overdrive suppression model by focal CoChR pacing of spontaneously active hearts at different frequencies. The optogenetic stimuli only changed beat rate and LOI position when the stimulus frequency was faster than the spontaneous frequency. Furthermore, once pacing stopped, the LOI immediately returned to its previous rhythm and location, with no apparent memory of pacing (Fig. S11). This mechanism is consistent with prior suggestions based on experiments in dissociated embryonic cardiomyocytes (*3, 28*), but to our knowledge has not previously been identified in the intact embryonic heart.

We then asked whether a spatially extended model of an excitable-to-oscillatory transition could account for the above observations on LOI position and response to perturbations. We simulated resistively coupled Morris-Lecar oscillators undergoing SNIC bifurcation, with a linear spatial gradient of input current, *I*, to model the observed clustering of the LOI in the anterior-left quadrant. Despite dynamic current noise and static heterogeneity of excitability, the simulations produced coherent waves which emerged from a single LOI and swept the tissue under a broad range of parameters (Supplementary Text, Fig. S12). A time-dependent offset in the current recapitulated the beat-rate statistics, as in the zero-dimensional QIF model (Fig. S12, Fig. 2, F and G). Thus, our model can explain how noisy and gradually changing single-cell excitability properties can produce abrupt and tissue-wide changes in action potential dynamics.

Fig. 4J summarizes the spatiotemporal dynamics of electrical activity in the earliest vertebrate heartbeats. The heart becomes electrically coupled and excitable before the first spontaneous beats, so the very first beat triggers a full-fledged action potential that coherently sweeps the tissue. Overdrive suppression among many spontaneously active units then assures the presence of one, and only one, locus of initiation in a single beat. This biophysical determination of the early LOI is not driven by a molecular “pacemaker” specification program, but rather emerges by the interaction of many noisy oscillators. Isl1+/Nkx2.5-pacemaker cells likely begin to act as the LOI later in development.

Although the morphogenesis and molecular details of early heart development have been thoroughly studied, the mechanisms underlying the onset of physiological function have been little explored. One could imagine many intermediate activity states between quiescence and coherent, periodic, tissue-wide beating. The mapping of gene expression onto electrical activity is complex, non-linear, and non-local. Due to gap junction couplings, the electrical properties of cells can change dramatically between *in vivo* and explanted conditions. All-optical electrophysiology allowed us to characterize the evolution of the earliest heartbeats *in situ* as development progressed. Our data show that the spatial structure undergoes a rapid step transition to tissue-scale engagement, but the temporal structure goes through a noisy intermediate between quiescence and periodicity. Remarkably, a simple dynamical model captured all of these features. We argue that any molecularly detailed model of heartbeat initiation must recapitulate the basic SNIC bifurcation dynamics described here. We speculate that these dynamical mechanisms ensure a stable heartbeat despite variable single-cell trajectories toward bioelectrical maturity.

## Acknowledgments

We thank C.E. and C.G. Burns for providing *Nkx2*.*5:ZsYellow* transgenic fish; I. Scott for providing the zACNE20 enhancer element; Y. Shen and R.A. Campbell for providing FR-GECO1c plasmid; H.C. Davis and F.P. Brooks for developing microscope control software; J. Miller-Henderson, K. Hurley, and A.R. Murphy for fish husbandry; and members of the Megason lab for assistance with fish husbandry.

## Author contributions

A.E.C, B.Z.J., and S.G.M. conceived the project and designed experiments. B.Z.J. performed all experiments, data analysis, and numerical modeling. B.Z.J. constructed molecular tools and transgenic fish lines. B.Z.J. wrote all new code used in analysis and simulations. Y.Q., J.D.W.C., and B.Z.J. performed optical engineering of microscopes. B.Z.J., A.E.C. and S.G.M. wrote the manuscript with input from all authors. A.E.C. and S.G.M. supervised the project.

## Data and materials availability

Data are available upon reasonable request to the corresponding authors.

## Supplementary Information

### Methods

#### Zebrafish strains and maintenance

All vertebrate experiments were approved by the Institutional Animal Care and Use Committee of Harvard University or the Harvard Medical Area Standing Committee on Animals. The zebrafish (*Danio rerio*) AB wild-type strain was used for all experiments and generation of transgenic fish. Adult fish were raised at 28.5 °C on a 14 h light/10 h dark cycle. Embryos were collected by crossing female and male adults (3 – 24 months old). The transgenic lines used in this study are: TgBAC(−36*nkx2*.*5*:*zsYellow*) (*1*), Tg(*zACNE20:2xLyn-mCherry*) (this study), Tg(*zACNE20:CoChR-eGFP-P2A-FRGECO1c*) (this study), Tg(*zACNE20-myl7:CoChR-eGFP-P2A-FRGECO1c*) (this study).

#### Morpholinos, RNA constructs, and microinjection

Morpholinos *tnnt2a-*MO (4 ng) (*2*), *cacna1c*-MO1 (2 ng), *cacna1c*-MO2 (2 ng) (*3*) (Gene Tools) were injected at the one-cell stage. Synthetic mRNAs were transcribed from linearized pMTB plasmid (*4*) containing insert of interest using the mMessage mMachine SP6 *in vitro* transcription kit (Thermo Fisher Scientific) and injected at the one-cell stage with 0.1% phenol red (Table S1).

#### Sample preparation for live functional imaging

Embryos were injected at the 1-cell stage with 20 pg mRNA encoding alpha-bungarotoxin, a peptide acetylcholine receptor blocker, to immobilize tail movements (*5*), 4 ng *tnnt2a* morpholino if performing spatially resolved recordings to eliminate cardiac motion artifacts, and the appropriate mRNAs for reporters and channelrhodopsins (Table S1). Embryos were raised at 28.5 °C to 16-20 somite stage (17-20 hpf). Egg chorions were removed by immersing embryos in 1 mg/mL Pronase protease (Sigma) in 0.3x Danieau buffer (17.4 mM NaCl, 0.21 mM KCl, 0.12 mM MgSO_4_, 0.18 mM Ca(NO_3_)_2_, 1.5mM HEPES, pH 7.2) for 7 minutes at room temperature followed by gentle mechanical agitation. A mount was cast in 1.5% agarose in 0.3x Danieau using a custom “shark-fin” mold (acrylic “fine detail plastic”, Shapeways; Fig. S1) in 35 mm or 100 mm Petri dishes. Embryos were mounted in 0.3x Danieau and affixed by placing a 0.17 mm glass coverslip over the mount array (25x imaging) or with a 0.15% low-melt agarose overlay solution (2x imaging).

#### Functional imaging with patterned optogenetic stimulus

Live imaging was performed at 28.5 °C or at room temperature using custom optical setups (Fig. 1A, (*6, 7*)). For recordings of jGCaMP7f dynamics, widefield illumination was supplied (0.5 – 1 mW/mm^2^) using a 488 nm laser (Coherent Obis) or a 470 nm LED. For spectrally orthogonal stimulation and recording, a 594 nm LED or laser (Cobolt Mambo 100 mW) was used for recording (1 – 2 mW/mm^2^ when recording at 50 Hz, 0.6 mW/mm^2^ when recording at 10 Hz), and blue light from a 488 nm laser or 470 nm LED (0.5–25 mW/mm^2^) was patterned by programming a digital micromirror device (ViALUX DLP7000 or Texas Instruments Lightcrafter DLP3000). Imaging was performed through a 25x NA 1.0 objective (Olympus XLPLN25XSVMP2), or 2x NA 0.5 objective (Olympus MVPLAPO 2XC). Data were recorded on an Orca Flash v4.0 camera (Hamamatsu) at 10-100 Hz framerate. Temporal control of lasers, DMD, and acquisition was performed using custom software through a National Instruments DAQ interface.

#### Confocal microscopy and morphological analysis

Live imaging was performed at 28.5 °C using a LSM 980 confocal microscope (Zeiss). All image stacks were filtered with a 5×5×1×1 (XYZT, pixel x pixel x section x frame) median filter to reduce shot noise before further processing. Approximate cell sizes and location of the inner edge of the heart cone were determined by manual tracing of maximum intensity projection images in ImageJ. Intensity and distance measurements were performed with Python scripts using the Scipy stack.

Cellular-resolution imaging of the first heartbeats was performed by scanning a single plane at 3 Hz. Images were filtered with a 5×5×1×1 (XYZT, pixel x pixel x section x frame) median filter and 2x downsampled in space. Active area was calculated in each frame by summing the number of pixels with intensity greater than 45% of a reference peak value (99.9^th^ percentile of all intensities in the video). To generate videos (Movie S5), the original recording was filtered with a Gaussian kernel ([σ_x_, σ_y_, σ_t_] = [2, 2, 3]), and ΔF/F was obtained by pixelwise dividing each intensity by the 10^th^ percentile over time.

#### Time series analysis for embryo array experiments

The full analysis pipeline is detailed in the Supplementary Text. Briefly, raw videos were denoised by detrending the mean of the entire field of view and correcting pixel-wise for photobleaching using the method described in (*6*). Frames with crosstalk from blue stimulus were replaced by the previous frame. Individual heart regions of interest (ROIs) in the array were segmented by identifying pixels with a normalized power within a frequency band of 0.1 Hz – 2.5 Hz above a manually set threshold (typically 50 – 60%). ROIs were refined by calculating the Pearson correlation of each pixel to the mean of the initial ROI. A new ROI was defined from pixels whose Pearson correlation was above a threshold. The time series spatially averaged over each refined ROI was extracted and converted into a ΔF/F value. Spikes were identified using the *scipy* function “peak_detect”. The analysis was performed on 2-minute video blocks that each contained multiple embryos.

Recordings of individual hearts with the 25x objectives were preprocessed as above but without image mean detrending, because the active heart comprised a large portion of the entire field of view. Spike- and stimulus-triggered averages were generated by using “peak_detect” and averaging a manually selected window around the transient maximum or light pulse. Further denoising was performed on the averages using a sub-Nyquist action potential timing (SNAPT) algorithm (*6, 8*). Activation maps were generated as described in Ref. (*9*).

The first heartbeat was detected in individual embryos by identifying the first instance where two consecutive inter-spike intervals were less than 120 s, which avoided misidentification of slow calcium transients or particle motion in the overlay solution as the first heartbeat. Frequency and coefficient of variation for individual embryos were extracted over 3-minute windows with a 90% overlap (steps of 18 s between points).

#### Locus of initiation (LOI) identification

The locus of initiation was identified from spike-triggered average videos of calcium indicator activity recorded at 50 – 100 Hz (full pipeline in Supplementary Text). Videos of the same embryo at different timepoints were registered to each other by choosing 2-D translation values that minimized squared error between *nkx2*.*5:ZsYellow* images taken immediately before each video. In gtACR2 targeting experiments, spontaneous activation maps were used to define an LOI target. In addition to this target several other ROIs were tested for each embryo. Because electrical coupling strength was unknown, we defined “Not LOI” targets as those which were above the 50^th^ percentile (47 μm) of distances (across all experiments) from the spontaneous LOI.

#### ODE and PDE simulations

Ordinary differential equation (ODE) and partial differential equation (PDE) simulations were performed in MATLAB and Python using Euler integration. The full analysis is detailed in the Supplementary Text.

#### Statistical analysis and plotting

Statistical analysis was performed using the *statannotations* and *scipy* Python packages. Plots were generated in Matplotlib. Specific tests and notation are listed in figure captions.

## Supplementary Figures

**Table S1.** Figure-wise detail of genotypes, treatments, and imaging parameters. *zACNE20-myl7:CoChR-eGFP-P2A-FRGECO1c* transgenics were supplemented with FR-GECO1c mRNA because the combination of experimental timing close to onset of expression and low absolute brightness of the protein resulted in low (but detectable) signal-to-noise ratio in the pure transgenics.

| Figures | Genotype | Treatment | Imaging parameters |
| --- | --- | --- | --- |
| 1, 2B – I, S5, S6, S7C, S8D – E, Movie S1 | AB | 20 pg αBTX mRNA, 75 pg jGCaMP7f mRNA | NA 0.5, 2x objective, 10 Hz widefield |
| 2A, S3, Movie S5 | Tg(*zACNE20:2xLyn-mCherry*) (+/-) | 20 pg αBTX mRNA, 75 pg jGCaMP7f mRNA, 4 ng *tnnt2a*-MO | NA 1.0, 20x water objective, #1 coverslip, 3 Hz confocal |
| 3B – D, Movie S2 | AB | 20 pg αBTX mRNA, 100 pg FR-GECO1c mRNA, 20 pg CoChR-eGFP mRNA | NA 0.5, 2x objective, 10 Hz widefield |
| 3E – F, S11, Movie S6 | Tg(*zACNE20-myl7:CoChR-eGFP-P2A-FRGECO1c*) (+/-) | 20 pg αBTX mRNA, 4 ng *tnnt2a*-MO, 100 pg FR-GECO1c mRNA* | NA 1.05, 25x water objective, #1 coverslip, 50 Hz widefield |
| 4A – C | TgBAC(-36nkx2.5:*ZsYellow*) (+/-, +/+) | 20 pg αBTX mRNA, 75 pg jGCaMP7f mRNA, 4 ng *tnnt2a*-MO | NA 1.05, 25x water objective, #1 coverslip, 100 Hz widefield |
| Movie S3 – S4 | AB | 20 pg αBTX mRNA, 75 pg jGCaMP7f mRNA, 4 ng *tnnt2a*-MO | NA 1.05, 25x water objective, #1 coverslip, 100 Hz widefield |
| 4E – I, Movie S7 | TgBAC(-36nkx2.5:*ZsYellow*) (+/-, +/+) | 20 pg αBTX mRNA, 4 ng *tnnt2a*-MO, 100 pg FR-GECO1c mRNA, 50 pg gtACR2 mRNA | NA 1.05, 25x water objective, #1 coverslip, 50 Hz widefield |
| S2 | TgBAC(-36nkx2.5:*ZsYellow*) (+/-, +/+) | 20 pg αBTX mRNA, 75 pg jGCaMP7f mRNA | NA 0.5, 2x objective, 10 Hz widefield |
| S4 | TgBAC(-36nkx2.5:*ZsYellow*) x Tg(*zACNE20:2xLyn-mCherry*) | 20 pg αBTX mRNA, 4 ng *tnnt2a*-MO | NA 1.0, 20x water objective, #1 coverslip, confocal z-stack |
| S9 | TgBAC(-36nkx2.5:*ZsYellow*) x Tg(*zACNE20:2xLyn-mCherry*); Tg(*zACNE20-myl7:CoChR-eGFP-P2A-FRGECO1c*) x Tg(*zACNE20:2xLyn-mCherry*); Tg(*zACNE20-myl7:CoChR-eGFP-P2A-FRGECO1c*) (+/-); Tg(*zACNE20:CoChR-eGFP-P2A-FRGECO1c*) (+/-) | 20 pg αBTX mRNA, 4 ng *tnnt2a*-MO | NA 1.0, 20x water objective, #1 coverslip, confocal z-stack; NA 1.05, 25x water objective, #1 coverslip, 100 ms exposure snap |
| S10 | AB | 20 pg αBTX mRNA, 75 pg jGCaMP7f mRNA, 4 ng *tnnt2a*-MO; 20 pg αBTX mRNA, 75 pg jGCaMP7f mRNA, 2 ng *cacna1c*-MO1, 2ng *cacna1c*-MO2 | NA 0.5, 2x objective, 10 Hz widefield |

**Table S2.** Simulation parameters for the Morris-Lecar model under different bifurcations. The model is specified in Supplementary Eq. 1.

|  | SNIC | Saddle-node | Supercritical Hopf | Subcritical Hopf |
| --- | --- | --- | --- | --- |
| C | 1 | 1 | 1 | 1 |
| $E_L, E_{Ca}, E_K$ | -80, 60, -90 | -80, 60, -90 | -78, 60, -90 | -78, 60, -90 |
| $g_L, g_{Ca}, g_K$ | 8, 20, 10 | 8, 20, 10 | 8, 22, 10 | 1, 4, 4 |
| $m_h, n_h$ | -20, -25 | -20, -25 | -23.4, -45 | -30, -45 |
| $k_m, k_n$ | 15, 5 | 15, 5 | 12.826, 5 | 7, 5 |
| $\tau$ | 1 | 0.159 | 1 | 1 |

**Figure S1.**
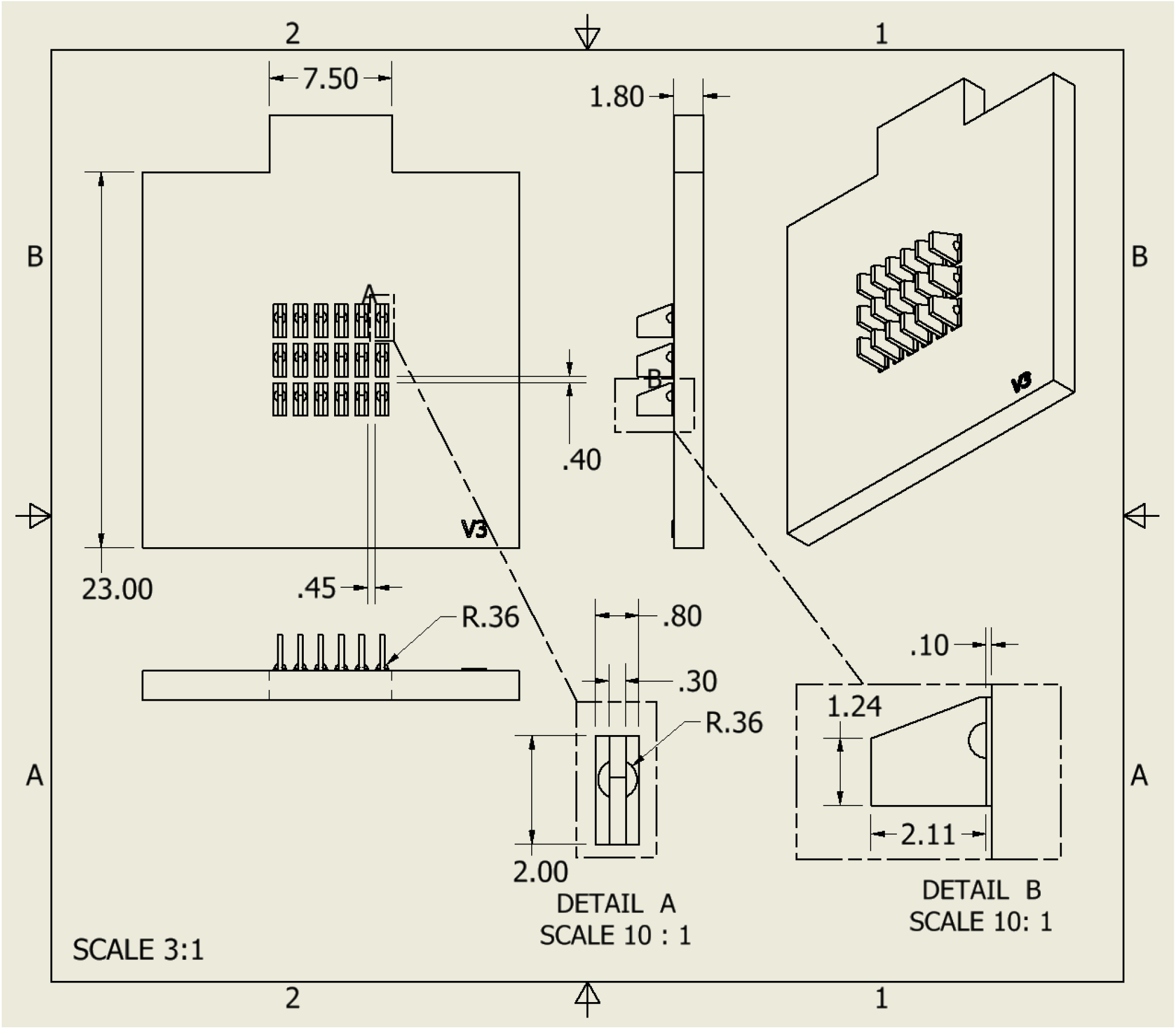
Engineering diagram for manufacture of shark-fin mount. All measurements in millimeters. Fits inside a 35 mm dish, accommodates a 22 mm coverslip.

**Figure S2.**
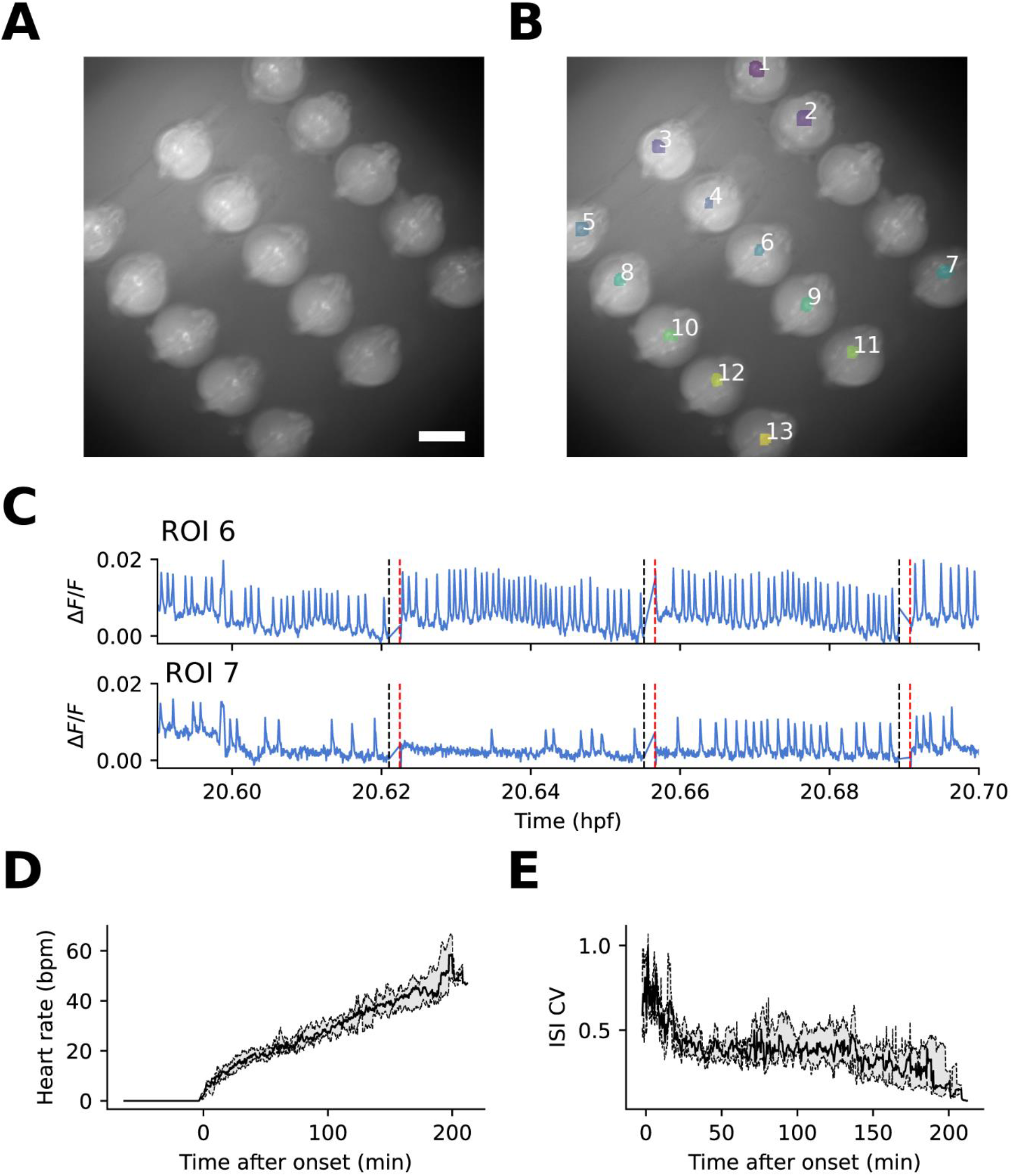
Calcium dynamics colocalize with nkx2.5 expression. (**A**) Tg(−36*nkx2*.*5:ZsYellow*) expression at 19.5 hpf. (**B**) Locations of first heartbeats segmented from FR-GECO1c transients overlaid onto (A). (**C**) Example ΔF/F traces of early cardiac activity reported by FR-GECO1c. Black and red lines indicate seams between recordings. (**D – E**) Statistical moments of early heartbeat dynamics reported by FR-GECO1c. Scale bar 500 μm. (D – E) Median ± IQR, N = 10 embryos.

**Figure S3.**
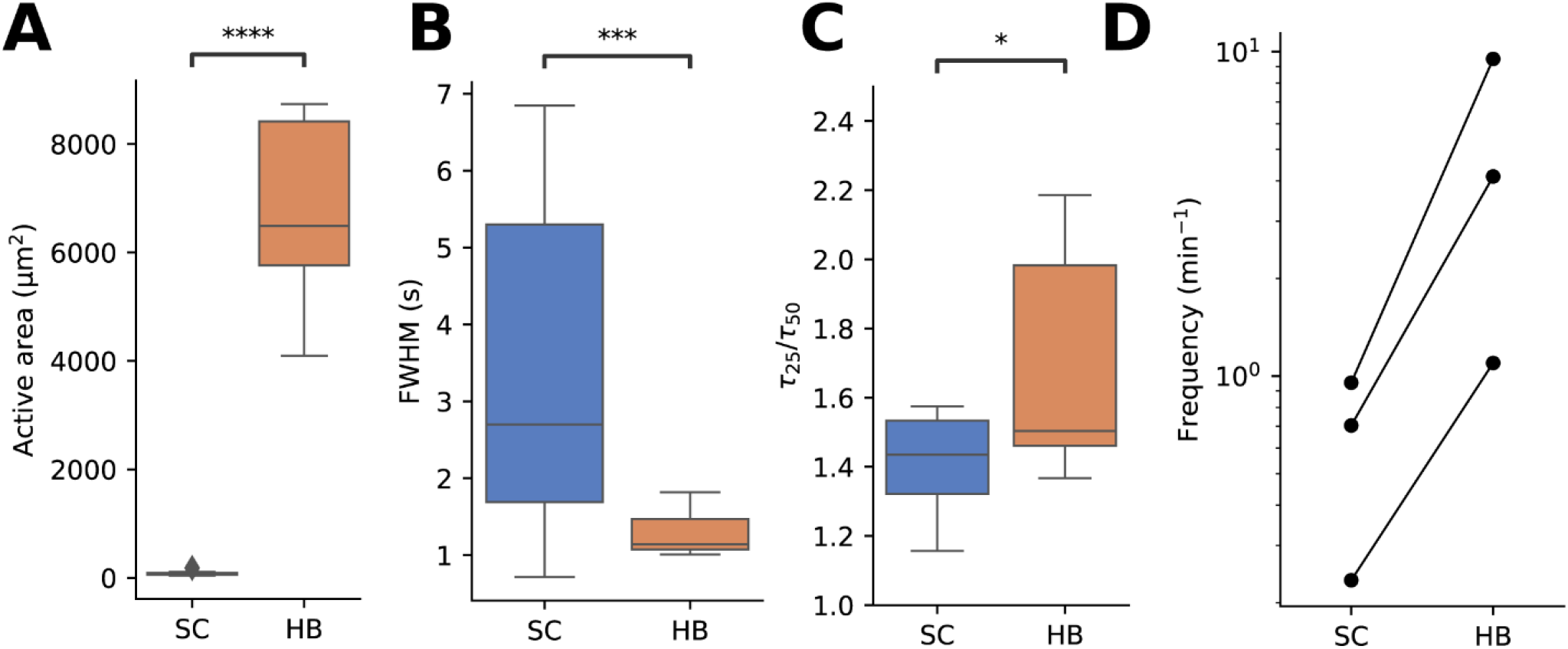
Dynamics of single-cell transients observed before the first heartbeat. (**A**) Active area of single-cell (SC) transients (74 ± 32 µm^2^) was much lower than that of the first heartbeats (HB, 7000 ± 1400 µm^2^). \*\*\*\**p* = 6.5e-11. (**B**) Single-cell transients were longer lived (3.4 ± 2.1 s) than the first heartbeats (1.3 ± 0.2 s). FWHM = full-width half-maximum. \*\*\**p* = 1.3e-4. (**C**) Single-cell transients had a longer plateau phase than the first heartbeats, as quantified by the ratio of peak width at 25% (τ_25_) and 50% (τ_50_) of maximum height. \**p* = 0.015. (**D**) Single-cell transients were rare (0.2 – 0.7 per minute) compared to the first heartbeats (1 – 9.5 per minute). Data from 19 single-cell transients and 50 beats across N = 3 embryos. Mean ± SD for all values in caption. p-values from two-sided Mann-Whitney test.

**Figure S4.**
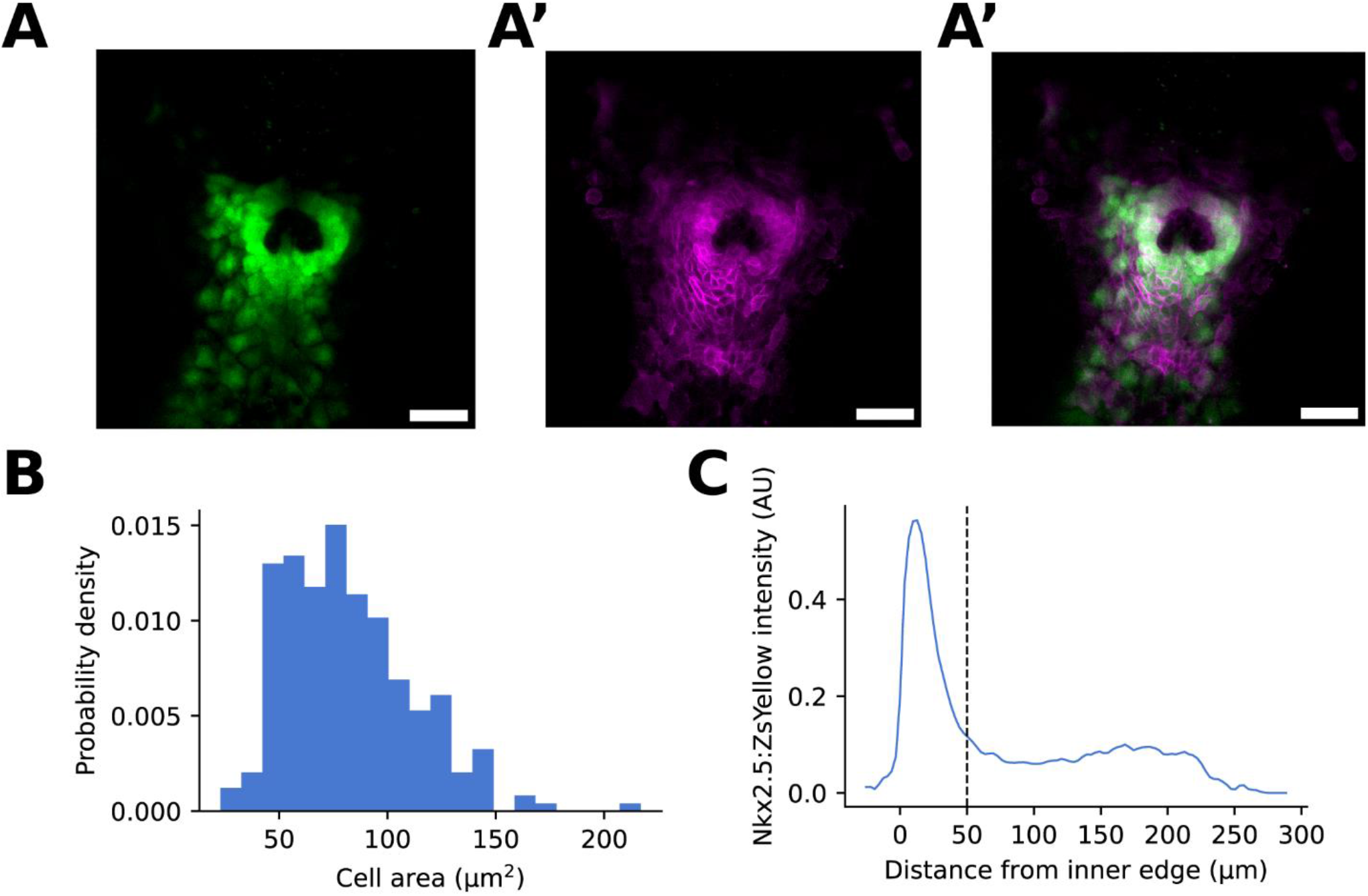
Tissue geometry of the heart cone. (**A – A’’**) Maximum intensity projection fluorescence images of 21-somite stage Tg(*-36nkx2*.*5:ZsYellow; zACNE20:2xLyn-mCherry*) heart cone. (A) *nkx2*.*5:ZsYellow*. (A’) *zACNE20:2xLyn-mCherry*. (A’’) Merge of (A) and (A’). Scale bars 50 µm. (**B**) Cell area distribution. Mean area 82 µm^2^ (± 29 µm^2^ SD). (**C**) Relative *nkx2*.*5:ZsYellow* intensity as a function of distance from the inner edge of the heart cone. A region approximately < 50 µm from the inner edge had stronger expression than more distal cells. N = 5 embryos, 254 cells in (B).

**Figure S5.**
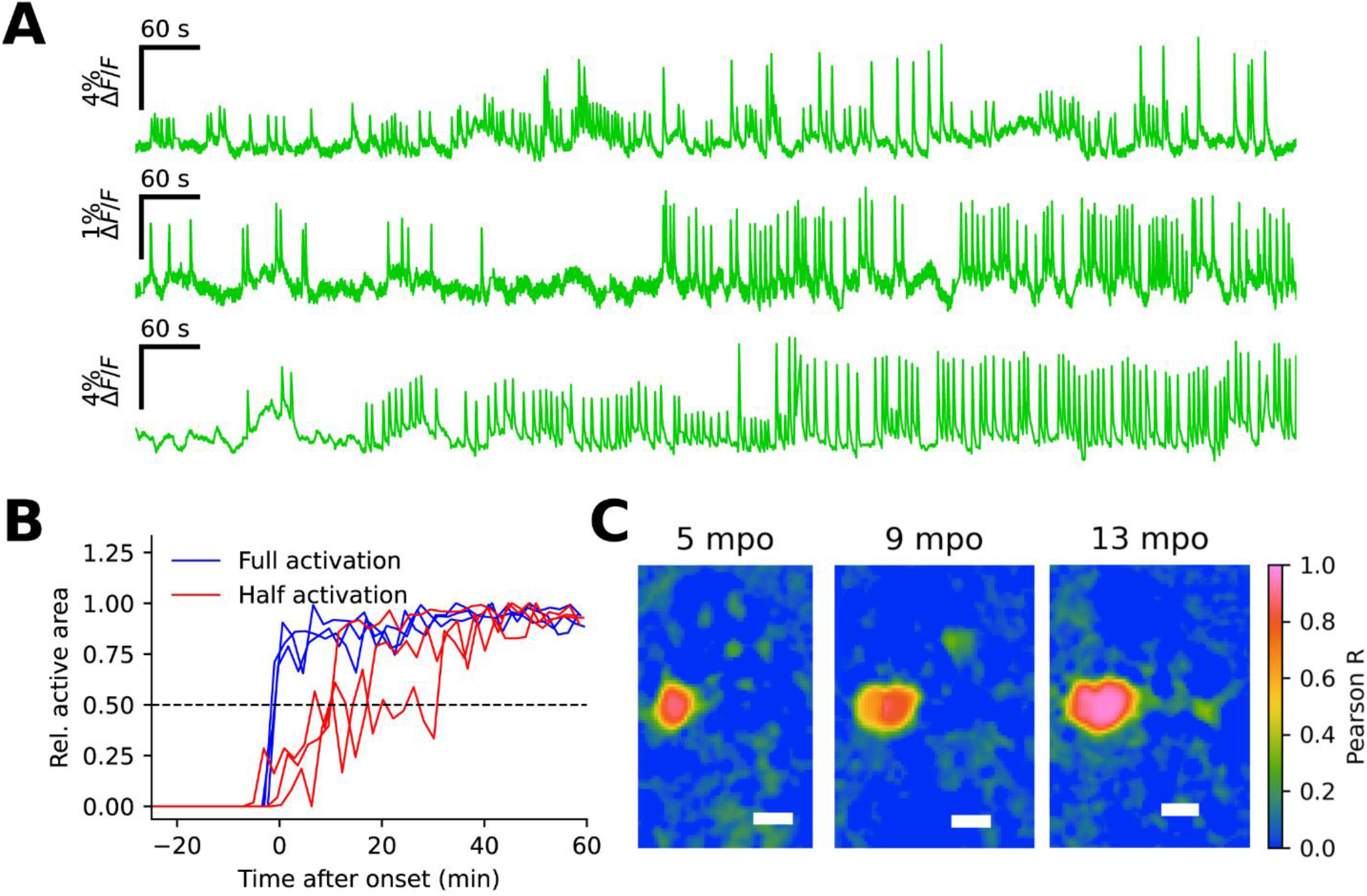
Rare initiation of spontaneous activity in half of the heart primordium. (**A**) jGCaMP7f ΔF/F traces starting from 2 minutes after the first heartbeat in embryos with spontaneous initiation in half the heart. The traces were averaged over the whole heart, so traces that engaged half the heart had approximately half the apparent amplitude of traces that engaged the whole heart. (**B**) Relative active areas of the traces in (A) compared to representative traces of embryos with full-tissue initiation. Here area was averaged over 2 min. (**C**) Pearson correlation of individual pixels with the mean segmented heartbeat jGCaMP7f activity at different timepoints. The sharp boundary in the 9 minutes post onset (mpo) image is due to roughly half the heart being engaged in some but not all beats. Same experiments as in Figure 1. Scale bars 100 µm.

**Figure S6.**
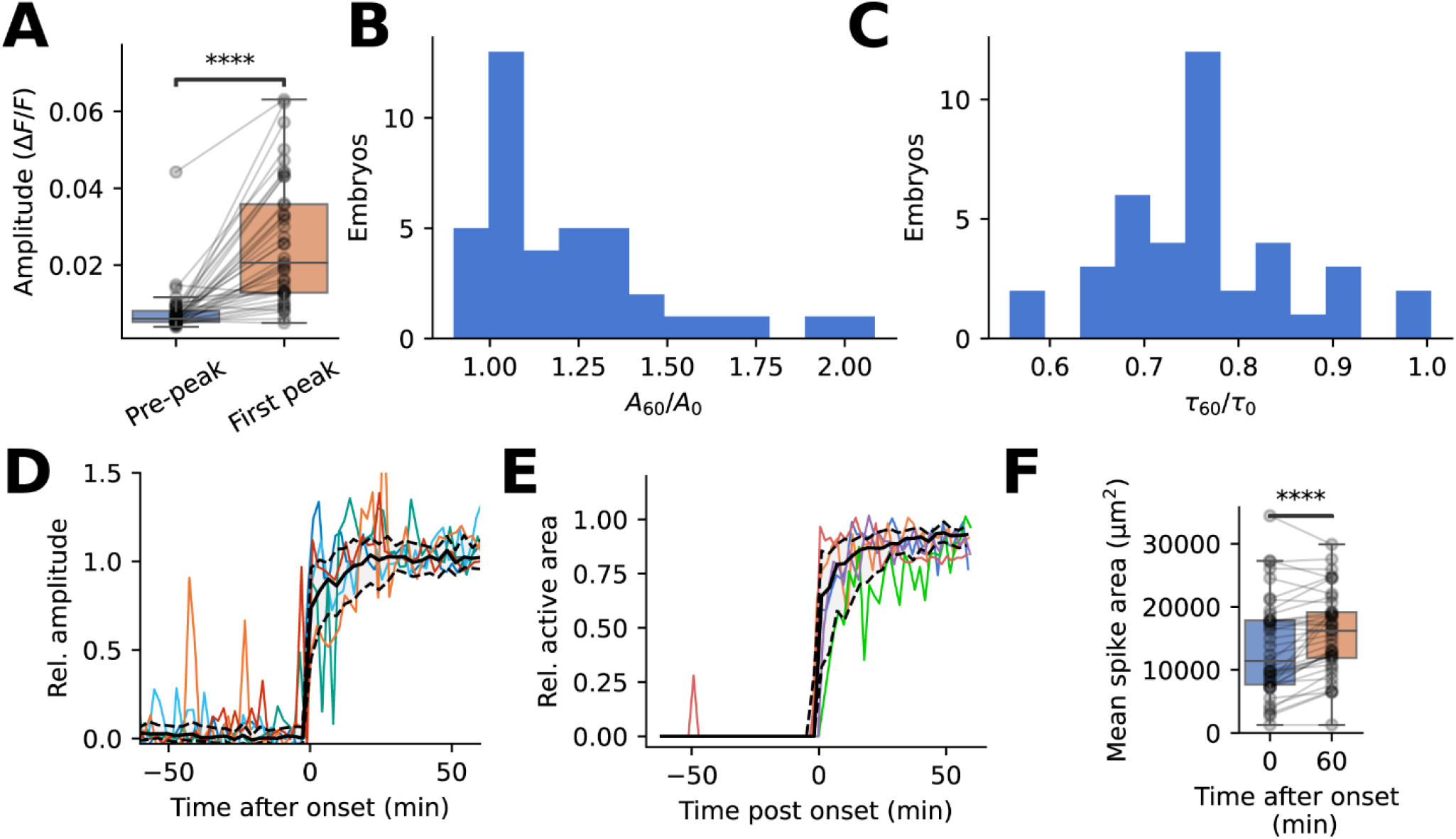
Development of calcium transients after onset of the heartbeat. (**A**) Amplitude of the first detected calcium transient compared to maximum fluctuation from mean in the previous 2 minutes. \*\*\*\**p* = 8.6e-9, paired t-test. (**B**) Ratio of calcium transient amplitudes (ΔF/F) 60 minutes after onset (*A*_60_) and 0 minutes after onset (*A*_0_). (**C**) Ratio of transient widths (full-width half-maximum) 60 minutes post onset (mpo) (*τ*_60_) and 0 mpo (*τ*_0_). (**D**) Amplitude of calcium transients as a function of time, aligned relative to first beat and normalized relative to initial beats. (**E**) Active of calcium transients as a function of time, aligned relative to first beat and normalized relative to initial beats (Methods). (D - E) Colored lines show representative single-embryo traces. Black line shows median. Shading and dashed lines show interquartile range. (**F**) Calcium spike area increased from 13490 ± 7730 µm^2^ at 0 mpo to 15840 ± 6400 µm^2^ at 60 mpo. **** p = 5.4e-5, paired t-test. (A – F) N = 39 embryos across 3 experiments, same individuals as Figures 1-2.

**Figure S7.**
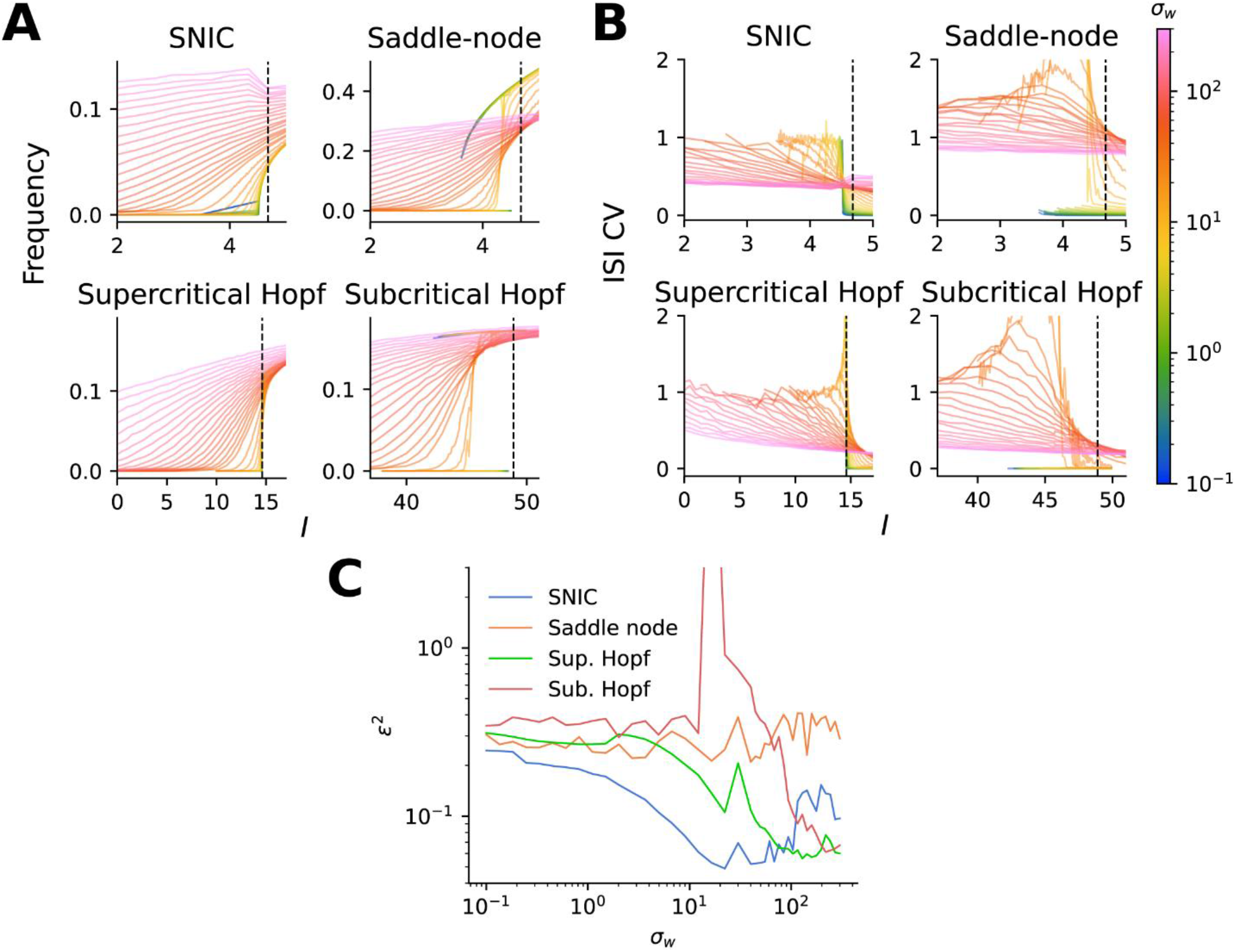
Statistical moments of spiking under codimension-1 bifurcations in the Morris-Lecar model. (**A**) Mean frequency (**B**) and inter-spike interval (ISI) coefficient of variation (CV) in simulated Morris-Lecar oscillators as a function of input current *I* and noise σ_w_. Parameters, given in Table S2, were selected to drive each of the four types of co-dimension 1 bifurcations. (**C**) Squared error (Supplementary Text) of experimental fits to each bifurcation at different values of *σ*_w_.

**Figure S8.**
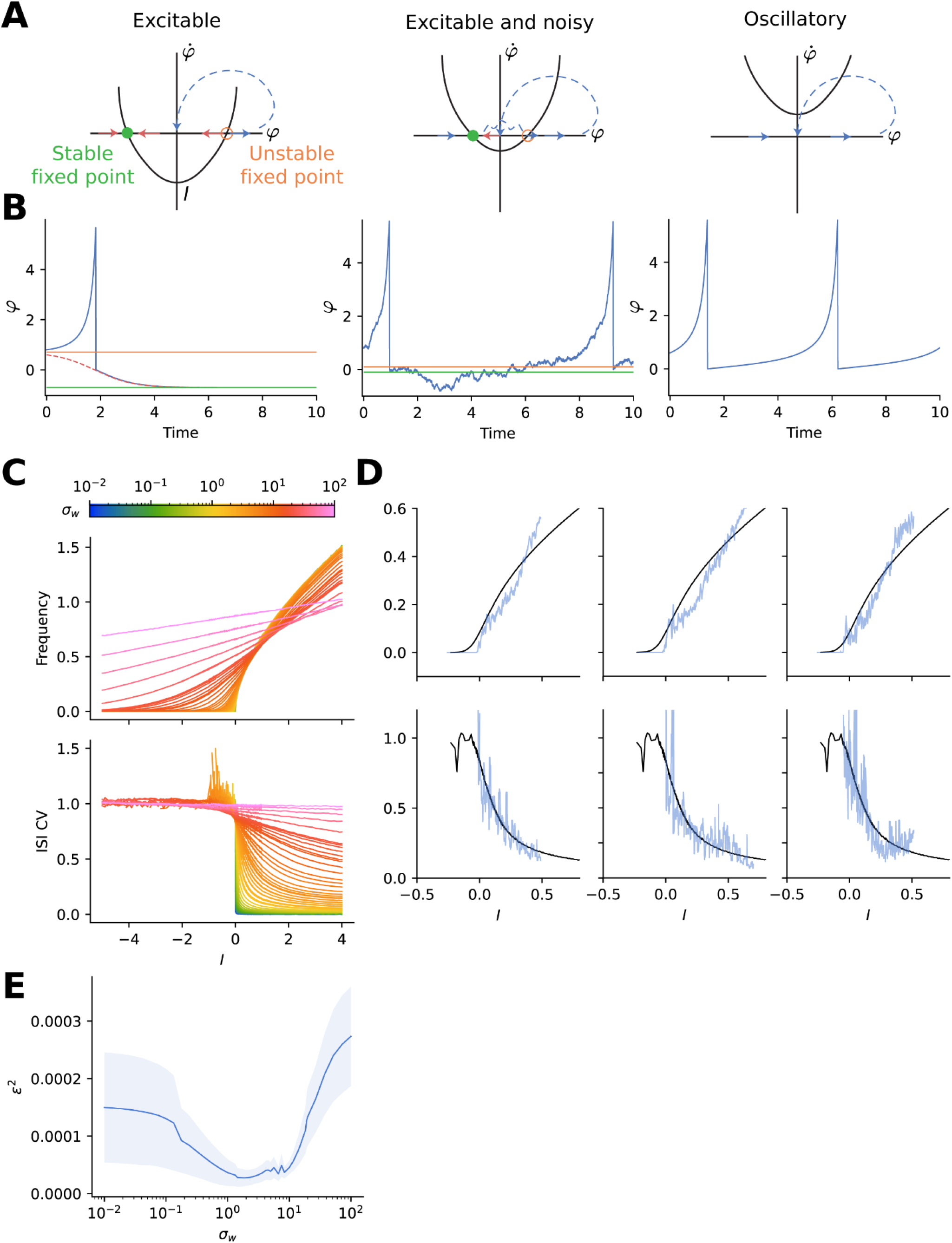
A noisy quadratic integrate-and-fire model captures the experimental statistics of the first heartbeats. (**A**) Spiking regimes across the SNIC bifurcation of the noisy quadratic integrate-and-fire (QIF) model, shown by 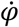 as a function of *φ*. (**B**) Simulated example traces for each spiking regime. (**C**) Simulated frequency and ISI CV for different values of noise σ_w_. (**D**) Example fits (top: frequency, bottom: CV) of individual embryo data (blue) to simulation (black) with choices of σ_w_ which minimized squared error. (**E**) Mean squared error of fits of simulated spike statistics to data as a function of σ_w_ (N = 39 embryos).

**Figure S9.**
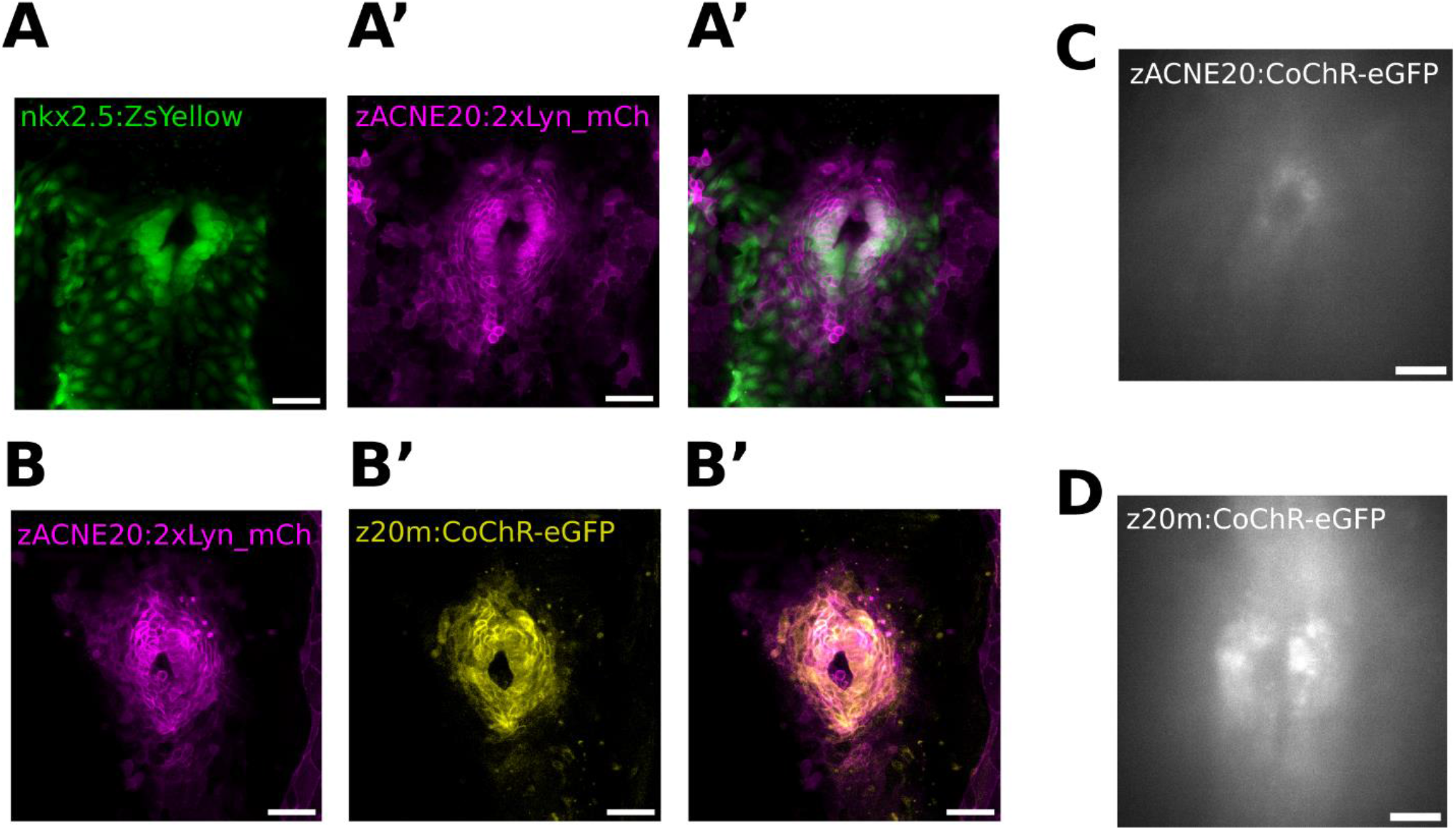
Expression patterns of *zACNE20* and *zACNE20-myl7* (*z20m*) promoters in the early heart. (**A-A”**) Maximum intensity projection images of the heart primordium in a 20-somite stage (ss) Tg(−36*nkx2*.*5:ZsYellow;zACNE20:2xLyn_mCherry*) embryo. (**B-B”**) Maximum intensity projection images of the heart cone in a 22 ss stage Tg(*zACNE20:2xLyn_mCherry; zACNE20-myl7:CoChR-eGFP-P2A-FRGECO1c*) embryo. (**C – D**) *z20m* produces stronger expression than *zACNE20* at the same stage. Widefield images at 20-21 ss.

**Figure S10.**
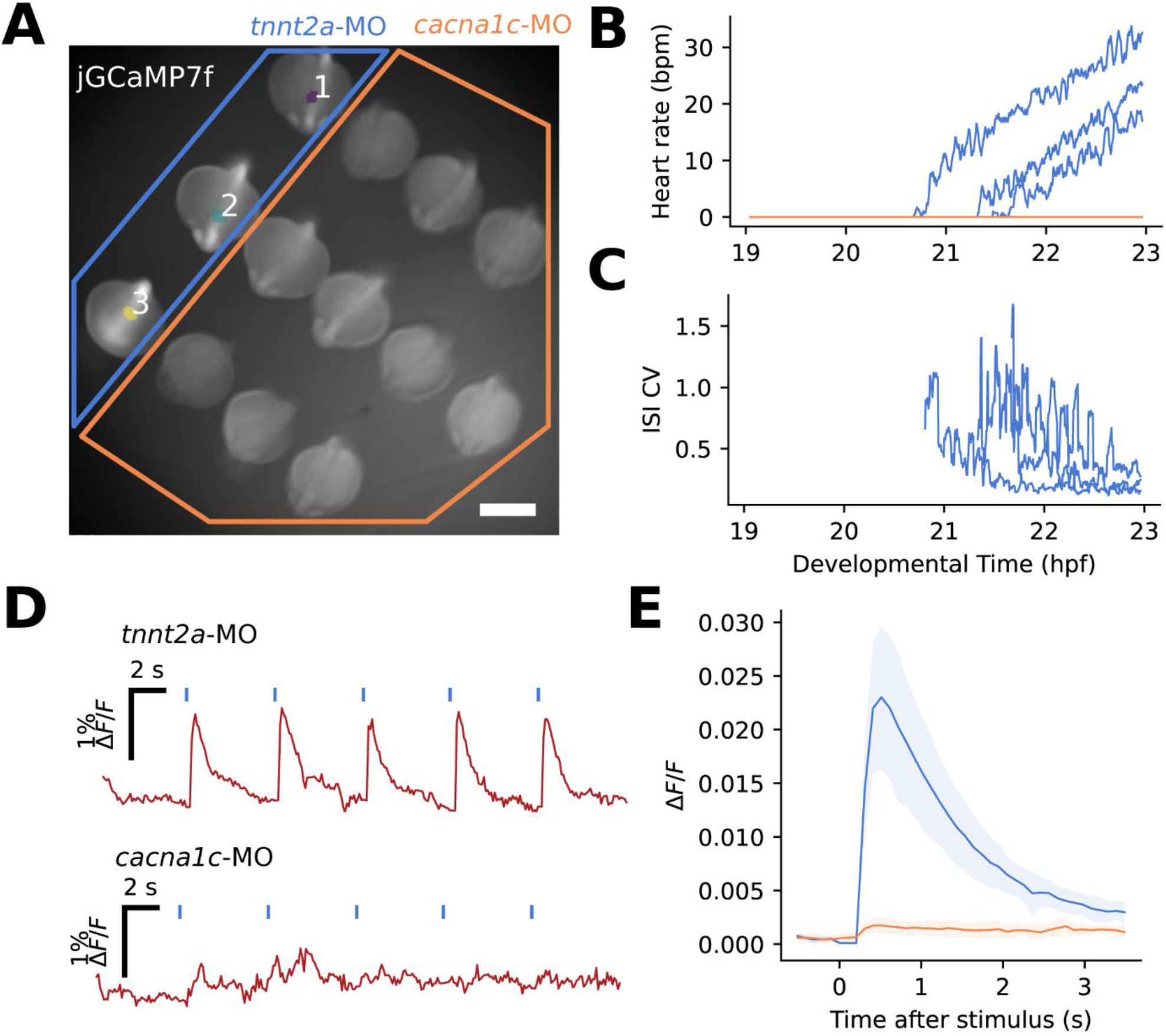
The L-type calcium channel is required for excitability and initiation of spontaneous activity. (**A**) Automatic segmentation (Methods) located heartbeats in control embryos but not in embryos injected with the *cacna1c* morpholino. (**B – C**) The heartbeats became faster and more regular after initiation in *tnnt2a*-MO embryos. Manually selected ROIs in the *cacna1c* morphants did not show spontaneous activity. (A – C) N = 3 embryos *tnnt2a*-MO, N = 10 embryos *cacna1c*-MO. All embryos were strain AB injected with jGCaMP7f mRNA. Scale bar 500 µm. (**D – E**) *cacna1c* morphants were not responsive to optogenetic stimulus. (D) Example traces of FR-GECO1c dynamics (red) with pulsed CoChR stimulus (blue) directed to the entire heart of individual embryos. (E) Stimulus-triggered average of calcium activity in control (blue) and *cacna1c* morphants (orange, population mean ± SD). (D – E) N = 11 embryos *tnnt2a*-MO, N = 14 embryos *cacna1c*-MO. All recordings acquired at 10 Hz. All embryos were *zACNE20-myl7:CoChR-eGFP-P2A-FRGECO1c* (+/-).

**Figure S11.**
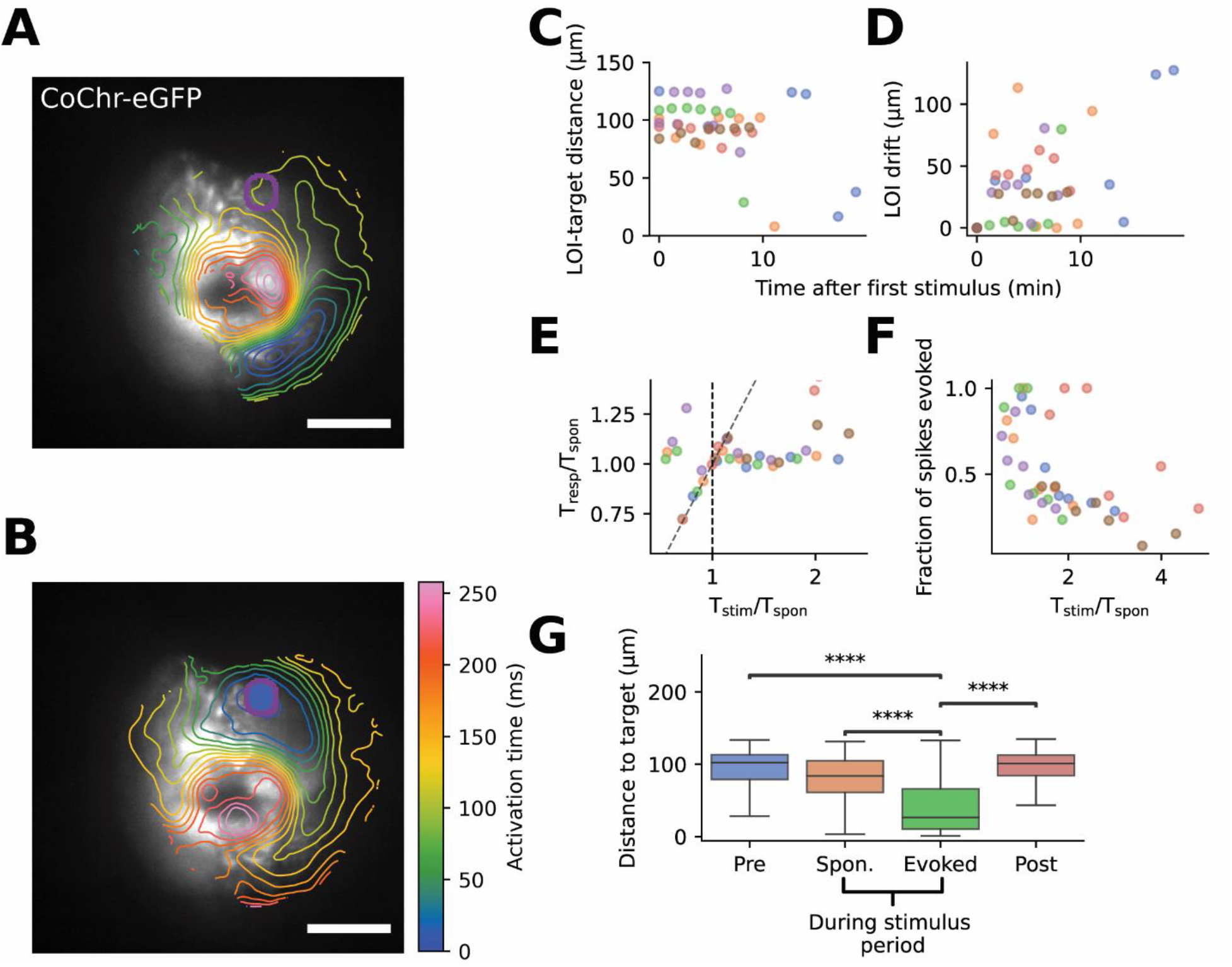
Optogenetic pacing induces overdrive suppression in the heart cone. (**A**) Activation map of spontaneous activity (spike-triggered average, n = 9 spikes). Purple circle indicates region to be stimulated in (B). (**B**) Activation map of evoked activity (spike-triggered average, n = 19 spikes) in the same heart as (A). Scale bars 50 μm. (**C**) Distance between spontaneous LOI and fixed “target region” after repeated pacing at different frequencies at the target region. (**D**) The drift of the spontaneous LOI from its first recorded position was uncorrelated with repeated pacing. (**E**) A target region away from the spontaneous LOI was paced with period *T*_stim_. Heartbeat was characterized by *T*_spon_ = period of spontaneous activity in absence of stimulus, and *T*_resp_ = period of activity during periodic stimulus. CoChR activation only paced the heart (i.e. *T*_resp_/*T*_stim_ ≈ 1, dotted line) when *T*_stim_ < *T*_spon_. At the highest stimulus frequencies, the showed a period doubling, i.e. *T*_resp_/*T*_stim_ ≈ 2. (**F**) Ratio of number of evoked beats to number of total beats (spontaneous + evoked) during the 30-second stimulus period. At a lower pacing interval, evoked beats comprised a larger fraction of the total. “Evoked beats” were defined as beats which occurred within 200 ms of a pulse of blue light. (C – F) Each color represents one fish. (**G**) Distance between the target region and the LOI of each individual beat grouped by timing and whether the beat was evoked. Evoked beats had LOI significantly closer to target than any other category. **** *p* < 4.3e-38, number of beats collected per category – “Pre”: 306; “During”: 309; “Stim”: 363; “Post”: 281. (C – G) *N* = 6 fish.

**Figure S12.**
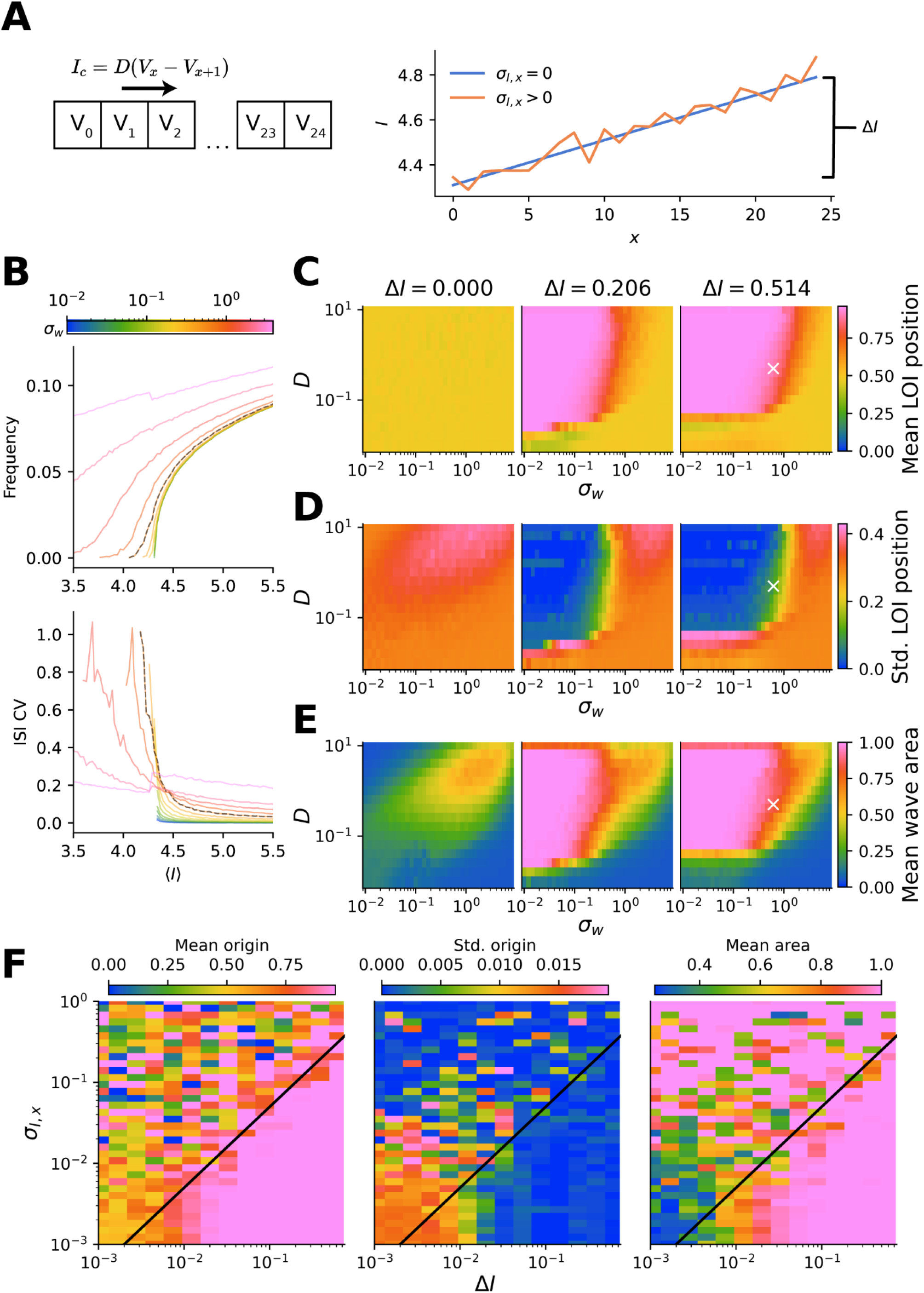
Robustness of wave geometry in heterogeneous coupled SNIC oscillators. (**A**) Schematic for chain of Morris-Lecar oscillators coupled by a current *I*_*c*_ which depends on coupling strength *D* (left); and spatial distribution of *I* for different choices of *ΔI* and *σ*_I,x_ (right). (**B**) Mean frequency (*f*) and inter-spike interval coefficient of variation (ISI CV) for a chain of 25 oscillators with *D* = 0.5, and current gradient *ΔI* =0.514. Simulation with *σ*_w_ = 0.618 (compare qualitatively to experiments) marked with dashed black lines. (**C – E**) Simulated wave properties as a function of *σ*_w_ and *D*. Here *I* attained its highest value at normalized *x* = 1.0. (C) Normalized mean LOI position. (D) Standard deviation of LOI position. (E) Normalized wave area. Parameter choice for dashed line in (B) indicated with white “X”. (**F**) Wave geometry as a function of *ΔI* and *σ*_I,x_. Black line indicates the line *σ*_I,x_ = 0.5*ΔI*. For sufficiently large *ΔI*, full-tissue wave propagation occurred with stable LOI at one end of the chain. In high variability, i.e. large *σ*_I,x_, a stable LOI driving full-tissue waves (low position standard deviation, high normalized mean area) can appear at positions that are not the end of the chain. (E – F) ⟨*I*⟩ = 4.68, *σ*_w_ = 0.03.

## Supplementary Movies

**Movie S1 – Parallel measurements of heartbeat in 18 embryos**. Embryos expressed jGCaMP7f and fluorescence was recorded at 10 Hz. Recording at approximately 22 hpf. Raw jGCaMP7f intensity in grayscale. ΔF/F in green. Scale bar 500 μm.

**Movie S2** – **Parallel all-optical electrophysiology in 15 embryos**. Embryos expressed CoChR and FR-GECO1c ubiquitously and red fluorescence was recorded at 10 Hz. Blue light stimuli were targeted to hearts but are indicated as offset blue dots (bright when illuminated) for clarity. Recording at approximately 22 hpf. Brightfield image in grayscale. Scale bar 500 μm.

**Movie S3 – Calcium dynamics before observable mechanical contractions**. Brightfield (left) and jGCaMP7f (right) videos taken consecutively at 10 Hz (20 **–** 21 hpf). Playback at 2X speed. Scale bar 50 μm.

**Movie S4 – Calcium dynamics with mechanical contractions**. Brightfield (left) and jGCaMP7f (right) videos taken consecutively at 10 Hz on the same embryo as Movie S3, 1 hour later. Playback at 2X speed. Scale bar 50 μm.

**Movie S5 –** The **first heartbeat at cellular resolution**. Two Tg(*zACNE20:2x:Lyn-mCherry*) (red) embryos expressing jGCaMP7f (cyan) imaged at ∼3 Hz. Videos are synchronized by the first detected heartbeat. Scale bar 50 μm.

**Movie S6 – Electrical wave propagation evoked by targeted CoChR activation before spontaneous cardiac activity**. Tg(*z20m:CoChR-eGFP-P2A-FRGECO1c*) embryo recorded at 50 Hz. CoChR-eGFP in grayscale. FR-GECO1c ΔF/F in red. Blue light stimulus in blue. Recording at approximately 19.5 hpf. Scale bar 50 μm.

**Movie S7 – Optogenetic hyperpolarization transiently perturbs wave geometry and locus of initiation**. Tg(*-36nkx2*.*5:ZsYellow*) embryo expressing gtACR2 and FR-GECO1c. Two consecutive 50 Hz recordings with different targets for optogenetic silencing. ZsYellow in grayscale. FR-GECO1c ΔF/F in red. Blue light stimulus in blue. Recording at approximately 21 hpf. Baselines set separately for blue-illuminated intervals (separated by white frames). Scale bar 50 μm.

## Supplementary Text

### Image analysis

#### (A) Preprocessing

Images were downsampled in space (4X along each axis). In optogenetic perturbation experiments, frames with optical crosstalk were identified by matching laser and camera triggers from NIDAQ waveforms and were replaced by the previous frame. In embryo array experiments, to correct for fluctuations in LED power or background light, the image-mean temporal fluctuations in intensity were subtracted from each pixel by linear regression. This step was skipped for high-magnification single-heart experiments because the heart comprised a large portion of the field of view and hence dominated the image-mean dynamics. Photobleaching dynamics were determined by taking the minimum intensity in a time window of 2 seconds around each frame (chosen to be longer than a typical heart calcium transient). In embryo array experiments, photobleaching correction performed on heart-average traces after segmentation. In single-heart experiments, the image mean was used. The signal was divided by the photobleaching trace.

#### (B) Heart segmentation in array experiments

Embryos were identified in arrays using local thresholding of mean intensity (Python *skimage*). Fast Fourier transforms in time were performed on each pixel. The power spectrum of each pixel was taken and normalized to the total power (area under curve). The fraction of power contained in the range (0.1Hz, 2.5Hz) was calculated. This fraction was smoothed in space using a 5×5 median filter, and a binary mask was generated by taking pixels with the power fraction above a threshold that was manually determined for each experiment. A 3×3 binary opening was performed on this mask, and connected ROIs were identified.

The ROIs identified from above were taken as initial guesses of heart locations. The mean trace in the time domain was taken for each ROI, and pixel-wise cross-correlation (lag 0) in the time domain was performed on pixels within the corresponding segmented embryos. A binary mask was generated from pixels with Pearson correlation coefficient exceeding a threshold that was manually determined for each experiment. 5×5 binary opening followed by 3×3 binary dilation were performed on this mask, and ROIs were identified as hearts. When measuring the spatial extent of active calcium dynamics, the last morphological operations were skipped to prevent overestimation of size. For measuring spatial extent of active calcium dynamics, the threshold for correlation coefficient was set to 0.65.

For 2-3 hour recordings, the above segmentation was performed on 2-minute blocks. The ROIs were then linked together in time starting from the last time point. For a given time point *t*, each detected ROI was assigned to the ROI in time point *t*+1 with the closest centroid, provided the distance was less than 15 pixels. ROIs in *t*+1 that were not successfully matched were then propagated to *t*. ROIs in *t* that were not successfully matched were then given new labels. The unlinked ROI video was then compared to the linked ROI video, and ROIs that were present in less than 15% of the unlinked frames were discarded. All ROIs with size less than 100 pixels were then replaced with the ROI nearest in time with size above this threshold. When measuring the spatial extent of active calcium dynamics, missing timepoints were not filled in.

#### (C) Spike detection

ΔF/F was determined by dividing the fluorescence signal by its 10^th^ percentile in time. Peaks were detected using *scipy peak_detect* using minimum heights, prominences, and width ranges that were determined manually for each experiment.

#### (D) Activation maps

Peak detection was performed on the mean trace of the brightest 30% of each image, and spike-triggered average (STA) videos were obtained. A B-spline representation of the STA of each pixel was obtained using scipy.interpolate.splrep (t=15 knots, s=0.05). The activation time of each pixel was determined by finding the time with the fastest upstroke of ΔF/F, 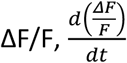. This was found by identifying the positive-to-negative crossing of 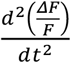 closest to the half-maximum of ΔF/F and preceding the maximum value of ΔF/F. Signal-to-noise ratio (SNR) was estimated by dividing the amplitude of ΔF/F at each pixel by the standard deviation of the residual of the spline smoothing. A SNR threshold for each recording was set as the lower of 10 (manually chosen) or a value automatically generated by the triangle threshold algorithm over the pixel-wise SNR distribution for the entire image. Pixels with SNR below the threshold were discarded. 3×3 morphological opening followed by 3×3 closing was used to define a “valid region” for the activation map. Activation times of discarded pixels were replaced with the mean activation time of valid pixels in a 13×13 neighborhood. Times were 3×3 median filtered and then Gaussian filtered with σ=1. Finally, a quadratic fit of all pixels in a 26×26 neighborhood with an activation time difference less than 350 ms was performed as in (*9*).

### Supporting models

Although excitable-to-oscillatory transitions have been widely proposed to have functional significance in neurons and other biological systems, there have been few observations of these phenomena without experimental intervention, and especially *in vivo*. We studied bifurcations of excitable systems to explain the temporal statistics of heartbeat initiation and the onset of propagating waves. We used three models: A) the Morris-Lecar model (*10*); B) the quadratic integrate-and-fire (QIF) model (*11*); and C) a spatially extended system of diffusively coupled Morris-Lecar oscillators. These models demonstrate respectively: A) that the early heart rate and variability corresponds to a saddle-node on invariant-circle (SNIC) bifurcation; B) the explanatory power of qualitative bifurcation types irrespective of the specific model; and C) an explanation of full-tissue oscillatory dynamics with varying locus of initiation (LOI).

#### (A) Comparison of spike distribution moments under different bifurcations using the Morris-Lecar model

Given the modulation of a single system parameter (codimension-1), there are only four bifurcations by which a stable equilibrium can disappear, and four bifurcations by which a stable limit cycle can appear (*12*). These are as follows.

Stable equilibria can disappear or lose stability via:

- Saddle-node bifurcation
- Saddle-node on invariant circle (SNIC) bifurcation
- Supercritical Hopf bifurcation
- Subcritical Hopf bifurcation Stable limit cycles can appear or disappear via:
- SNIC bifurcation (simultaneous with loss of stable equilibrium)
- Supercritical Hopf bifurcation (simultaneous with loss of stable equilibrium)
- Fold limit cycle bifurcation (often preceding a subcritical Hopf bifurcation)
- Saddle homoclinic orbit bifurcation (often preceding a saddle-node bifurcation)

The important properties of oscillatory dynamics which differ between these cases are the amplitude of oscillations, the ability to fire at arbitrarily low frequencies, and the presence of bistability of spiking and resting states. A detailed discussion can be found in (*12, 13*), and reference can also be made to Type I and Type II neural excitability originally observed experimentally by Hodgkin (*14*). We sought to describe the spontaneous transition from excitable to spiking (as opposed to forced transition from a stable fixed point to a co-existing limit cycle), so we considered the bifurcations that describe loss of stability in equilibria.

The Morris-Lecar (ML) model was originally developed to describe the contractions of the giant barnacle muscle fiber and achieves good correspondence to experimental data while only accounting for voltage-dependent calcium and potassium currents (*10*). It has since been widely used to study the spiking dynamics of excitable cells. The ML model can display all codimension-1 bifurcations in *I*, given appropriate choices of the other model parameters. The formulation we used is as follows (adapted from (*12*)):

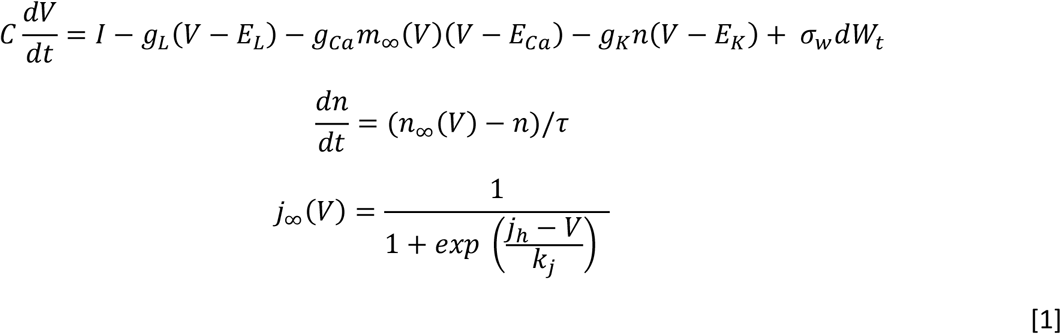

Where *C* is membrane capacitance, *V* is membrane potential, *I* is input current, and *g*_*i*_ and *E*_*i*_ are conductances where *i* is one of the leak (*L*), calcium (*Ca*), or potassium (*K*) currents (note that these physiological labels are arbitrary and the model does not require specific potassium or calcium channels). *m* and *n* are the gating variables for calcium and potassium respectively, and *τ* is the characteristic timescale of potassium gating. *J*_*∞*_ describes the steady-state gating function with a half-maximum at *j*_*h*_ and a slope factor of *k*_*j*_, where *j* ∈ {*m, n*}. We assume that calcium gating dynamics are fast compared to potassium gating and voltage dynamics, and by timescale separation arrive at a two-variable system. An additional noise term is added to capture the stochastic behavior of the data: *dW*_t_ is a Wiener process step and *σ*_w_ is its standard deviation. When combined with the rest of the dynamics, this means that subthreshold trajectories approach an Ornstein-Uhlenbeck process when far from the spike threshold.

Here we are interested in the spike statistics under noise in the regimes of each bifurcation. This problem has previously been analyzed, contrasting general categories of neural type I and type II excitability (*15*–*17*). We explored spiking dynamics under noisy forcing with physiologically realistic parameter sets which would generate each possible codimension-1 bifurcation in the absence of noise (Table S1, (*12*)). We numerically solved these equations using Euler integration with *Δt* = 0.002. As initial conditions we numerically solved for the fixed points of the system and added a small random increment in *V* and *n*. In cases with more than one fixed point (saddle-node bifurcations), we chose an unstable one. We allowed the simulation to run until *t* = 1000 to exclude transient dynamics before collecting spike statistics over the following time window of *T* = 50 000. We plotted frequency *f* and coefficient of variation (CV; *λ*) for different values of I and 0.01 < *σ*_w_ < 300.

To compare the experimental data to the simulations, we performed the following linear transformations:

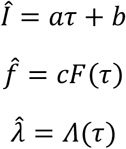

Where *τ* is the real time after onset of the heartbeat, *F*(*τ*) is the experimental beat frequency, *⋀*(*τ*) is the experimental CV, and *a, b, c* represent free parameters of time scaling, time offset, and frequency scaling respectively. Since *λ* is already a dimensionless quantity, it requires no scaling.

For each value of *σ*_w_, we jointly fit the mean and CV of experimental data for individual embryos to minimize the mean squared error of all observations τ_i_ over values of *a, b, c*:

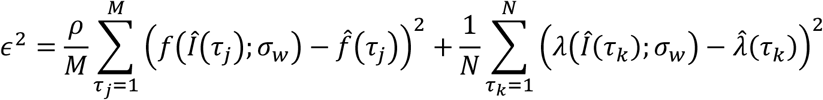

The functions *f*(*Î*) and *λ*(*Î*) were obtained by linearly interpolating the simulated values. *ρ* is an additional scaling factor that is set manually to enforce an equal weighting between individual residuals of *f* and *λ* and to create comparable error sizes between models that generate different absolute frequencies. Experimental observations of frequency and CV at times τ_i_ were considered separately (τ_j_ and τ_k_) to reflect the fact that a frequency of 0 contains information about the dynamics but does not have a defined CV. The individual residuals were normalized by the number of observations per moment to give equal weighting to the fits of CV and frequency. The fitting was performed using the AMPGO nonlinear optimization method (*18*), with manually tuned inequality constraints on *a, b, c*.

The *f*-*I* and *⋀*-*I* curves showed large differences between the bifurcations. Only the spike statistics in the SNIC bifurcation resembled the data, as shown above (Figs. S5, S6). Oscillators near all other bifurcations showed a comparatively rapid transition to a non-zero frequency, followed by relative insensitivity of the frequency to *I* (Type II excitability). In contrast, oscillation frequency near a SNIC bifurcation was dependent on a slow portion of the trajectory near the saddle-node point, and its residence time in this neighborhood was strongly dependent on *I*. Thus, the SNIC bifurcation can have arbitrarily low frequency. Furthermore, the other bifurcations displayed *⋀* ≫ 1 near the bifurcation point, which is consistent with bursting spikes and was rarely observed in our data.

Quantitatively, the SNIC bifurcation resulted in the best fit to the data. For 1 < *σ*_w_ < 50, the SNIC parameters resulted in lower error than the other bifurcations (Fig. S7C). At the optimal value of *σ*_w_ = 22, the ε^2^ was 2-fold to 18-fold smaller in the SNIC bifurcation than the others and attained a minimum over all bifurcations and values of *σ*_w_ tested. Other types of bifurcation dynamics, such as bursting, have been observed experimentally in other electrically active cells (*19*). For *σ*_w_ > 50, the models became very similar to each other, because the stochastic component dominated the deterministic subthreshold dynamics of the system. Therefore, the spiking frequency approached a first passage time problem of the Wiener process in all cases.

#### (B) Noisy QIF model

The differential equation used for simulation is as follows, adapted from (*11*):

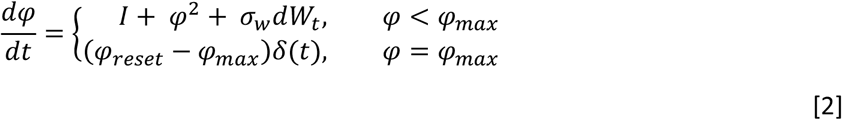

where *φ* is the dynamic variable, *I* is an excitability parameter (“input current”), *dW*_*t*_ is an increment of the Wiener process (uncorrelated noise sampled from a Gaussian distribution), and *σ*_*w*_ is the noise amplitude. Simulations and fitting were performed as in (A). In all simulations *φ*_*reset*_ = 0.

Briefly, a qualitative description of the dynamics is as follows. If *I* < 0 and 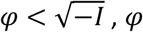, *φ* decays to resting value 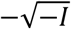 (Fig. S3A). If *I* < 0 and 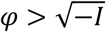, or *I* > 0, *φ* approaches infinity in a finite time. Practically, once *φ* reaches a finite value *φ*_*max*_, it is set back to *φ*_*reset*_, which we choose to be 0 (Fig. S3B). Therefore, if *I* < 0, the system is asymptotically stable. If *I* > 0, sustained oscillations will occur (Fig. S3B). Strictly speaking, neither excitation nor limit cycles are possible in 1-dimensional systems, but the case-wise definition of the differential equation allows qualitatively similar behavior to be described. Specifically, the QIF model is a normal form of the SNIC bifurcation, meaning all equations undergoing a SNIC bifurcation can be transformed onto it through an appropriate change of variables (*11*).

With *σ*_w_ > 0, spontaneous spiking can occur when *I* < 0 (Fig. S3C). This stretches out the increase in spike frequency *f*(*I; σ*_*w*_) as *I* is increased, and introduces a coefficient of variation (CV, *λ*(*I; σ*_*w*_)) in the inter-spike intervals (ISIs) (Fig. S3D). Given *σ*_w_ > 0, a “noisy but spontaneous” regime emerges. All three regimes (quiescent, noisy but spontaneous, and spontaneous) have clear qualitative correspondence to our data (Fig. 3, A – C). Quantitatively, ε^2^ was a function of *σ*_w_ with a global minimum, which also minimized the variation of ε^2^ across samples. These facts together demonstrate that the SNIC bifurcation is a good description of the experimentally observed dynamics and statistics.

#### (c) Formation of spatial structure in media of heterogeneous and noisy ML oscillators

To model a coupled medium of Morris-Lecar oscillators, we used the following equation:

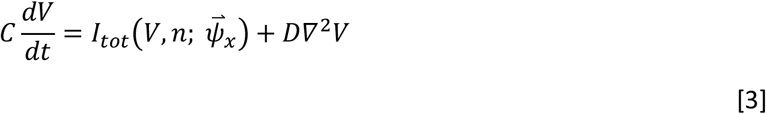

Where *I*_*tot*_ (*V, n;* 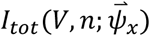) is the right-hand side of the first expression in [1] given all specified parameters 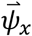 at position *x*, and *D* = *g*_*c*_*L*^2^, where *g*_*c*_ is the gap junction conductance and *L* is the length of a cell. All other equations are as in [1]. In numerical simulations, we used a 1-dimensional chain of 25 oscillators (approximately the number of cells observed from one end of the heart cone to the other), with a linear spatial gradient in *I* and all other parameters 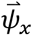 identical in each oscillator (Table S2). We detected spikes in each oscillator and identified propagating waves by grouping adjacent oscillators that fired within a specified interval of each other.

Our single-oscillator model showed that the increase in beat frequency during development could be recapitulated by an increase in the bifurcation parameter *I*. The anterior-left localization of the spontaneous LOI, and its mobility over time, suggested that the parameter *I* had both a variation across the heart, and likely some static noise. If the cardiac progenitor cells were not electrically coupled to each other, this scenario would cause different cells to beat at different frequencies, something we did not observe. We therefore explored the conditions under which the intercellular gap junctional coupling could cause multiple oscillators with different natural frequencies to phase lock.

We imposed a spatial gradient of *I* along the 25-unit chain using the SNIC parameters for the ML model. The mean-field relationships between *I* and frequency and ISI CV were similar to those observed in the single-oscillator model (Fig. S12A). To characterize the spatial structure of the simulations, we quantified the mean and standard deviation of the wave origin position, as well as the mean number of oscillators recruited by each wave (Fig. S12, B – D). We found that a small spatial gradient of *ΔI* = 0.514 (compared to a rate of change 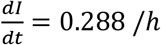 predicted by the single-oscillator model) was sufficient to induce consistent wave propagation across the medium from the end with higher *I*. This was true for large ranges of coupling strength *D* and noise strength σ_w_, including values whose mean field statistics approximated the experimental data (Fig. S12, A - D). Furthermore, small random deviations from the linear spatial gradient of *I* did not affect the position of the LOI or the extent of wave propagation (Fig. S12E). Larger deviations caused the LOI to move away from the end of the chain, but full-area waves occurred nonetheless (Fig. S12E).

A plausible biological interpretation of these results is that a graded tissue-scale signal which promotes electrical maturation of the cardiomyocytes underlies *ΔI*. Initially cells are relatively similar but have some intrinsic variability to their bioelectrical properties, resulting in a spatially variable LOI. The gradient sets a directional bias which grows over time, allowing the LOI to migrate to the anterior left as cells there increase their natural frequency. With sufficiently strong electrical coupling, the LOI can be maintained at one position on the short timescale (beat-to-beat) due to overdrive suppression, even as heartbeats pass through the entire tissue.

